# TMEM135 is an LXR-inducible regulator of peroxisomal metabolism

**DOI:** 10.1101/334979

**Authors:** Benjamin J. Renquist, Thushara W. Madanayake, Jon D. Hennebold, Susma Ghimire, Caroline E. Geisler, Yafei Xu, Randy L. Bogan

## Abstract

The liver x receptors (LXRs) are key regulators of systemic lipid metabolism. We determined whether transmembrane protein 135 (TMEM135) is an LXR target gene and its physiologic function. An LXR agonist increased TMEM135 mRNA and protein in human hepatocyte and macrophage cell lines, which was prevented by LXR knockdown. The human *TMEM135* promoter contains an LXR response element that bound the LXRs via EMSA and ChIP, and mediated LXR-induced transcription in reporter assays. Knockdown of TMEM135 in HepG2 cells caused triglyceride accumulation despite reduced lipogenic gene expression, indicating a potential role in β-oxidation. To determine physiologic importance, TMEM135 was knocked-down via siRNA in livers of fed and fasted C57BL/6 mice. Fasting increased hepatic fatty acid and NADH concentrations in control mice, consistent with increased fatty acid uptake and β-oxidation. However, in fasted TMEM135 knockdown mice, there was a further significant increase in hepatic fatty acid concentrations and a significant decrease in NADH, indicating an impairment in β-oxidation by peroxisomes and/or mitochondria. Conversely, hepatic ketones tended to increase in fasted TMEM135 knockdown compared to control mice, and because ketogenesis is exclusively dependent on mitochondrial β-oxidation, this indicates peroxisomal β-oxidation was impaired in knockdown mice. Localization studies demonstrated that TMEM135 co-localized with peroxisomes but not mitochondria. Mechanistically, proteomic and Western blot analyses indicated that TMEM135 regulates concentrations of matrix enzymes within peroxisomes. In conclusion, *TMEM135* is a novel LXR target gene in humans that mediates peroxisomal metabolism, and thus TMEM135 may be a therapeutic target for metabolic disorders associated with peroxisome dysfunction.

## Introduction

The liver x receptors (LXR) α (NR1H3) and β (NR1H2) belong to the nuclear receptor superfamily and are key regulators of cholesterol and fatty acid metabolism [1]. Oxysterols are natural LXR ligands [2] that increase during cholesterol loading. The LXRs heterodimerize with the retinoic acid receptor (RXR) and bind to LXR response elements (LXRE) within the promoters of target genes [2]. The LXREs belong to the direct repeat 4 (DR4) class that consist of two hexanucleotide half-sites separated by a 4-nucleotide spacer [3,4]. The protein products of known LXR target genes affect systemic lipid metabolism. Well known examples include increasing cholesterol efflux via ATP-binding cassette (ABC) sub-family A, member 1 (ABCA1) [5], and sub-family G, member 1 (ABCG1) [6]; limiting cholesterol uptake via induction of myosin regulatory light chain interacting protein (MYLIP) that targets the low-density lipoprotein receptor (LDLR) for ubiquitin-mediated degradation [7]; and increased lipogenesis via induction of sterol regulatory element binding protein 1c (SREBP1c, encoded by sterol regulatory element binding transcription factor 1 gene or *SREBF1*) [8,9]. Also, the *NR1H3* isoform of the LXRs is itself an LXR target gene [10,11], leading to auto-amplification of LXR actions in certain species and tissues. Although many LXR target genes have been discovered, additional target genes may remain unidentified.

We have previously investigated LXR actions within the steroidogenic corpus luteum of the ovary [12–15]. To uncover novel LXR target genes, a microarray experiment of rhesus macaque luteal cells treated in the presence or absence of a synthetic LXR agonist was performed. One gene that was differentially expressed in this analysis and not previously determined to be an LXR target gene was transmembrane protein 135 (*TMEM135*). Therefore, the objective of the current study is to determine if *TMEM135* is an LXR target gene and to identify its physiologic role in lipid metabolism.

## Materials and methods

### Microarray analysis

Procedures involving rhesus macaques were approved by the Oregon National Primate Research Center’s IACUC. Corpora lutea (CL) were collected from normally menstruating rhesus macaques 12 days after the midcycle luteinizing hormone surge, and mixed luteal cells were isolated and cultured from CL as described previously [16]. Luteal cell cultures were treated for 24 hours with 0.1% (*v:v*) DMSO or 1 µM T0901317 (T09) (n = 3 paired biologic replicates). Cells were harvested, mRNA were isolated and used for microarray analysis, and microarray data were analyzed as described elsewhere [17]. The criteria for differential expression of probes was a minimum 2-fold change in expression between T09 and DMSO treatments, p < 0.05 (paired t-test) with Benjamini and Hochberg correction for false discovery rate. Microarray data have been deposited in the National Center for Biotechnology Information’s Gene Expression Omnibus under accession number GSE130902.

### Cell lines

All cell lines were obtained from American Type Culture Collection (ATCC) and were cultured at 37°C, 5% CO_2_ in air, in a humidified incubator. Human cells (all derived from males) included the hepatocyte lines HepG2 (ATCC^®^ HB-8065^™^) and Hep 3B2.1-7 [Hep3B] (ATCC^®^ HB-8064^™^), and the monocyte line THP-1 (ATCC^®^ TIB-202^™^). Murine cells included the hepatocyte line (embryonic-derived) BNL 1NG A.2 (ATCC^®^ TIB-76^™^) and the macrophage line (male) RAW 264.7 (ATCC^®^ TIB-71^™^). Growth media for all cell lines except THP-1 was DMEM/F12 (Sigma Aldrich, Inc.) supplemented with Pen/Strep (100 units/ml penicillin and 100 µg/ml streptomycin) and 10% fetal bovine serum (FBS). THP-1 cells were incubated in RPMI media (Sigma) supplemented with 2-mercaptoethanol (0.05 mM), Pen/Strep, and 10% FBS. THP-1 cells were differentiated into macrophages by treatment with 12-myristate-13-acetate (PMA, 100 ng/ml) for 5 days prior to experimentation.

### Mice

All procedures involving mice were approved by the University of Arizona IACUC. Male C57BL/6 mice, 12 weeks of age, were used. Mice were group housed until initiation of experimental procedures, at which point they were switched to individual caging.

### Cell treatments

T0901317 (Cayman Chemical, Inc.) was dissolved in DMSO and used at a 1 μM concentration unless indicated otherwise. Cycloheximide (ACROS Organics, Inc.) was dissolved in DMSO and used at a final concentration of 50 μg/ml. The final DMSO concentration was held constant in all groups at less than 0.2% (*v:v*). For siRNA knockdown experiments, Silencer^®^ select pre-designed siRNAs (Life Technologies, Inc.) against human *NR1H2* (s14684), *NR1H3* (s19568), *TMEM135* (s35201), and a negative control siRNA (catalog 4390843) were purchased and transfected into HepG2 cells using Lipofectamine^®^ RNAiMAX (Life Technologies) or TransIT-siQuest^®^ reagent (Mirus Bio, LLC.) according to manufacturer instructions. All experiments were repeated 4-5 times on different days or with different passages of cells.

### Electrophoretic mobility-shift analysis (EMSA)

The DNA sequences for human *NR1H2*, *NR1H3*, and RXR α (*RXRA*) were synthesized by Life Technologies and cloned into the pTarget™ mammalian expression vector (Promega Corporation) using XhoI and KpnI restriction enzyme sites. Inserts were confirmed by DNA sequence analysis. Recombinant NR1H2, NR1H3, and RXRA proteins were produced using the TNT^®^ Quick Coupled Transcription/Translation System (Promega). Fluorescent probes were synthesized by Integrated DNA Technologies and corresponded to the LXRE1 sequence with IRDye 700 attached to the 5’ end, and probes corresponding to the LXRE3 sequence were labelled with IRDye 800. Additionally, unlabeled DNA oligos corresponding to LXREs 1-3, as well as mutated LXREs 1-3, were synthesized (Fig S1B). The EMSA binding reactions were carried out for 30 min at room temperature in 10 mM Tris, 50 mM KCl, pH 7.5; 3.5 mM DTT, 0.25% Tween-20, 1 μg Poly (dI.dC), 5 nM fluorescent probe (1 nM each for LXRE1/LXRE3 competition experiment), and 1.5 μl of TNT lysate per protein (non-induced TNT lysate substituted in control reactions). Unlabeled competitor DNA was added prior to the addition of fluorescent probe at a 200-fold molar excess unless indicated otherwise. Following the incubation period, EMSA reactions were resolved on a 6% TBE gel (Life Technologies). Imaging and densitometry were performed with a Li-COR Odyssey CLx and Image Studio version 3.1 software, respectively.

### Reporter assays

The region from −2662 to −1 bp (relative to translation start site) of the human *TMEM135* promoter was synthesized by Life Technologies. Additionally, the same region of the human *TMEM135* promoter was synthesized except that point mutations were introduced into each of the three LXREs (Fig S1B). The normal and mutant *TMEM135* promoters were cloned into the pGL4.17 vector (Promega) using SacI and XhoI restriction enzyme sites. A unique PacI site was identified in the region between LXRE1 and LXRE2, and a unique EcoRI site was identified in the region between LXRE2 and LXRE3. These restriction sites were used to generate six more *TMEM135* promoter/pGL4.17 constructs so that all possible combinations of wild type and mutated LXREs were obtained. All constructs were verified using DNA sequence analysis. These reporter vectors were transfected into HepG2 cells along with a β-galactosidase control plasmid (Promega) to normalize transfection efficiency. In some transfections, *NR1H3*/pTarget and *RXRA*/pTarget constructs were co-transfected. Vectors were transfected in a ratio of 60:20:10:10 (luciferase:β-galactosidase:NR1H3:RXRA) using Lipofectamine^®^ 3000 according to manufacturer recommendations, with no-insert plasmids substituted in control transfections to keep DNA concentrations constant. Each vector combination was transfected in triplicate to HepG2 cells in 96 well plates, and cells were incubated with DNA/Lipofectamine complexes for 24 hours at which point media were changed and cells were treated for an additional 48 hours with DMSO or T09. Cell lysates were prepared using mammalian protein extraction reagent (Fisher), and the lysate was fractionated to quantify luciferase and β-galactosidase activity using luminescent detection kits (Fisher). The ratio of luciferase to β-galactosidase activity was calculated, and the fold-change in T09 vs DMSO treated cells was determined for each transfection.

### Chromatin immunoprecipitation (ChIP) analysis

ChIP was performed as we have described elsewhere with minor modifications [13]. After fixation, nuclei from HepG2 cells were isolated by incubating cell pellets in 500 μl of a hypotonic buffer for 15 min on ice (20 mM Tris-HCL, pH 7.4, 10 mM NaCl, 3 mM MgCL_2_, with protease and phosphatase inhibitor cocktail from Promega). Triton X-100 detergent was added to a 0.5% *v:v* final concentration to lyse the cells followed by centrifugation at 1200 × g for 10 min to pellet nuclei. The supernatant was removed, and the nuclei pellet washed once with 500 μl hypotonic buffer. Nuclei were sonicated in RIPA buffer to produce chromatin of optimal length (approximately 200-1000 base pairs). The quantity of DNA used in each ChIP was held constant within replicates (8 to 10 μg). Antibodies used for ChIP (3 μg per reaction) were mouse monoclonal IgG2a including: 1) anti-NR1H3 (clone PPZ0412, R&D Systems catalog PP-PPZ0412-00), 2) anti NR1H2 (clone K8917, R&D Systems catalog PP-K8917-00), and 3) non-specific control IgG2a (clone MOPC-173, Abcam catalog ab18413). Immunoprecipitated chromatin was analyzed by QPCR in a duplex reaction with primers and a Taqman probe specific for the region spanning LXRE3 on the *TMEM135* gene, and a non-specific locus was simultaneously quantified in the same reaction as a control for non-specific DNA carryover (Table S1). Serial dilutions of input chromatin were included to verify linear amplification and PCR efficiency, and Ct values from ChIP reactions and input controls were used to calculate the percentage of DNA recovered relative to the input. Data were normalized by calculating the ratio of the TMEM135 LXRE3 to the non-specific locus.

### Semi-quantitative real-time PCR (QPCR)

Cells or tissue were homogenized in Trizol reagent for isolation of mRNA and were further purified over RNeasy columns according to manufacturer recommendations (Life Technologies). Concentrations of RNA were quantified by spectrophotometry, and 0.2-1 μg RNA was treated with DNAse I and reverse transcribed into cDNA using the High Capacity cDNA Reverse Transcription Kit (Life Technologies). Primer and probe sequences used for QPCR are listed in Table S1. The QPCR assays were performed on a StepOne Plus Real-Time PCR system (Life Technologies) using TaqMan^®^ MGB probe or SYBR green detection. Relative mRNA abundance was determined by extrapolation of threshold (Ct) values from a standard curve of serial cDNA dilutions and normalized to the housekeeping gene mitochondrial ribosomal protein S10 (*MRPS10*).

### Cell proliferation assay

HepG2 cells were plated at 15,000 cells per well in 96-well plates and transfected with siRNAs immediately. At the appropriate post-transfection time, viable cells were quantified using a CyQuant^®^ Direct Cell Proliferation assay (Life Technologies) according to manufacturer recommendations, and fluorescence (excitation 495 nm, emission 527 nm) determined using a Synergy H1 plate reader (BioTek Instruments, Inc.) in area scanning mode to account for uneven cell distribution. The number of cells per well was estimated by extrapolation from a standard curve of increasing cell numbers that was generated at the time of plating.

### Flow cytometry analysis of cell cycle

Freshly harvested HepG2 cell pellets were fixed by slow addition of 1 ml ice-cold 70% ethanol while vortexing and incubated at −20°C overnight. Fixed cells were harvested by centrifugation at 850 × g for 5 min and resuspended in 1 ml DNA staining buffer (100 μg/ml RNAse A, 50 μg/ml propidium iodide, in PBS). Cells were incubated for 30 min at 37°C, placed on ice, and analyzed within one hour. A Becton Dickinson FACSCANTO II flow cytometer equipped with a 488 nm argon ion laser and a 585/42 bandpass filter was used for detection of bound propidium iodide. List mode data files consisting of 10,000 events were acquired and the percentage of cells in G0/G1, S, and G2/M stages was determined with CellQuest PRO software (BD Biosciences).

### HepG2 ATP quantification

At 48 hours post-siRNA transfection, HepG2 cells were switched to serum-free, glucose-free, sodium pyruvate-free DMEM media and incubated at 37°C for 4 hours. Cellular ATP concentrations were determined using a homogenous luminescent ATP detection kit (Abcam catalog ab113849). Additional HepG2 cells treated in parallel to those used for ATP determination were lysed in RIPA buffer and a BCA protein assay (Fisher) was performed, and ATP content was normalized to protein concentrations.

### Triglyceride quantification and neutral lipid staining

HepG2 cells or frozen mouse liver were sonicated in PBS containing protease and phosphatase inhibitors (Promega) in an ice-water bath using a Branson 250 digital sonifier programmed to cycle at a 10% amplitude with 5 seconds on followed by 20 seconds off, and a 30 second total sonication time. Triglycerides were extracted using an organic solvent procedure optimized for triglycerides [18]. Dried extracts were dissolved in assay buffer (50 mM potassium phosphate, pH 7.0, 0.01% Triton X-100). Extracts (50 μl) were transferred to a clear 96 well plate, and 250 μl of triglyceride detection reagent (Pointe Scientific, Inc., catalog T7532) was added. Triglyceride standards (Pointe Scientific) diluted in assay buffer were also run with each assay. Plates were incubated at 37°C for 30 minutes with vigorous shaking, and absorbance at 500 nm was determined with a Synergy H1 plate reader. The protein concentration in sonicated lysates was determined using a BCA protein assay, and triglyceride content was normalized to protein content. Neutral lipids were visualized in HepG2 cells using LipidTox™ red neutral lipid stain (Life Technologies) according to manufacturer recommendations, and counterstained with DAPI to visualize nuclei.

### In vivo siRNA delivery and tissue harvest

Ambion^®^ In Vivo pre-designed siRNA (Life Technologies) against *Tmem135* (s91285) or *in vivo* negative control #1 siRNA (catalog 4459405) were complexed with Invivofectamine^®^ 3.0 reagent (Life Technologies) according to manufacturer recommendations. The siRNA complexes were delivered via a 200 μl tail-vein injection at a final siRNA concentration of 1 mg/kg body weight. Mice were weighed prior to siRNA injection and again before euthanization. Mice were euthanized 4 days after siRNA injection at 4 hours after lights on, with or without a preceding 12 hour fast. Serum, liver, heart, adipose, and skeletal muscle were collected and snap frozen. Tissues were powdered while frozen and stored at −80° C until used.

### Serum analytes

Total cholesterol, HDL cholesterol, and triglyceride concentrations in serum were determined as we have described previously [19]. Commercial enzymatic assays were used to quantify serum concentrations of β-hydroxybutyrate (BHB) (Cayman Chemical, catalog 700190), non-esterified fatty acids (NEFA) (Catachem Inc., catalog C514-0A), glucose (Pointe Scientific, Inc., catalog G7521), and a commercial ELISA was used to quantify serum insulin (Mercodia AB, catalog 10-1247) according to manufacturer recommendations.

### Liver analytes

Commercial kits were used for the extraction and quantification of liver NAD+ and NADH (Abcam Inc., catalog ab176723), and glycogen (BioAssay Systems, catalog E2GN), according to manufacturer recommendations. Concentrations of ATP and NEFAs in liver were determined as we have described elsewhere [20]. To quantify BHB, powdered liver was sonicated in PBS, acidified with 10% (*w:v*) trichloroacetic acid, centrifuged to remove proteins, and the supernatant was recovered and neutralized. Concentrations of BHB in the extract were determined using the same enzymatic assay described for serum. Concentrations of cholic acid in liver were quantified using a competitive enzyme immunoassay according to manufacturer recommendations (Cell Biolabs, Inc., catalog MET-5007).

### Liver subcellular fractionation

Powdered liver (approximately 50 mg) was homogenized for 40 strokes in a glass dounce homogenizer with a tight-fitting pestle using 600 μl of cold fractionation buffer (10 mM Tris-HCl, pH 7.4, 1 mM EGTA, 200 mM sucrose, protease and phosphatase inhibitor cocktail) on ice. The homogenate was transferred to a new tube, and the dounce homogenizer was washed with an additional 600 μl of fractionation buffer and transferred to the same tube. The homogenate was centrifuged at 600 × g for 10 minutes at 4°C to pellet unbroken cells and nuclei, and the supernatant transferred to a new tube. This low speed spin was repeated once more. Mitochondria and peroxisomes were pelleted by centrifugation at 7,000 × g for 10 minutes, and the supernatant (cytosolic fraction) was transferred to a new tube. The pellet was washed with 300 μl fractionation buffer and pelleted as before. The pellet was resuspended in 500 μl of fractionation buffer and a 30 μl aliquot was pelleted and solubilized in RIPA buffer for a BCA protein assay. Following protein assay mitochondria/peroxisome fractions were partitioned into 30 μg (protein content) aliquots followed by centrifugation and removal of the supernatant, and the pellets were frozen at −80°C.

### Proteomic analyses

The mitochondria/peroxisome-enriched fraction from a subset of fed control and TMEM135 knockdown mice was solubilized in RIPA buffer, proteins were precipitated with acetone, and proteins were sent to the Arizona Proteomics Consortium for mass spectrometry identification of proteins. Proteins were digested with trypsin and equal quantities of protein (500 ng) were loaded. Samples were analyzed on a Thermo Q Exactive Plus Orbitrap mass spectrometer. Peptide identification in the resultant tandem mass spectra was performed using Proteome Discoverer Software version 1.3.0.339 scanning with the SEQUEST algorithm against the mouse proteome database (Mouse_unitprotkb_proteome_2016_0720_cont.fasta). Data analysis was performed using Scaffold version 4.8.1, with a peptide identification threshold of 95.0% and a minimum protein identification threshold of 99.9% and 2 unique peptides.

### Western blot

All primary antibodies were obtained from Abcam unless otherwise indicated. Catalog numbers and final concentrations or dilutions of each antibody were: rabbit polyclonal anti-ACAA1 (catalog ab154091, 1 μg/ml), rabbit monoclonal anti-ACOX1 (catalog ab184032, 1:2500 dilution), rabbit monoclonal anti-CAT (catalog ab209211, 1:2000 dilution), rabbit monoclonal anti-SCP2 (catalog ab140126, 1:2500 dilution), rabbit polyclonal anti-TMEM135 (catalog ab222237, 0.4 μg/ml), mouse monoclonal anti-COX4I1 (catalog 14744, 0.25 μg/ml), rabbit polyclonal anti-PMP70 (catalog ab3421, 0.5 μg/ml), and mouse monoclonal anti-TUBB (Sigma Aldrich catalog T8328, 0.5 μg/ml).

Proteins (12.5 μg/well for mouse liver, 40 μg/well for HepG2 cells) were resolved on 4-12% Bis-Tris gels (Life Technologies) and subsequently transferred to nitrocellulose membranes. The membranes were blocked with 5% non-fat dry milk (NFDM) (*w:v*) in TBS with 0.1% (*v:v*) Tween (TBST) for 1 hour at room temp. All primary antibodies were diluted in TBST with 1% NFDM and incubated with the membranes on a rocking platform at 4° C overnight. The membranes were washed 4x with TBST, and IRDye^®^ 680RD or 800CW-conjugated secondary antibodies (LI-COR Biosciences, Inc.) were diluted 1:4000 in TBST + 1% NFDM and incubated with membranes for 1 hour at room temp. The membranes were washed as before. Imaging and densitometry were performed with a Li-COR Odyssey CLx and Image Studio version 3.1 software, respectively.

### Subcellular localization of TMEM135

The *TMEM135* coding region was amplified from cDNA of HepG2 cells using Phusion DNA Polymerase (ThermoFisher Scientific), and subcloned into the pTarget™ mammalian expression vector (Promega Corporation) using XhoI and KpnI restriction enzyme sites. The enhanced green fluorescent protein (EGFP) sequence was amplified via PCR from a pcDNA3-EGFP plasmid that was a gift from Doug Golenbock (RRID:Addgene_13031). The EGFP sequence was cloned in-frame on the N-terminus of *TMEM135* in the pTarget vector using BamHI and XhoI restriction enzymes. Inserts were confirmed by DNA sequence analysis.

A plasmid expressing mCherry with a peroxisomal targeting signal (PTS1) (RRID:Addgene_54520), and a plasmid encoding mCherry with a mitochondrial targeting sequence (RRID:Addgene_55102), were gifts from Michael Davidson. HepG2 cells were plated in black, clear-bottom 96-well plates at 25,000 cells per well, and the following day plasmids were transfected using Trans-IT 2020^®^ reagent (Mirus Bio) according to manufacturer instructions. At 48 hours post-transfection nuclei were stained with 2 μg/ml Hoechst 33342 for 30 min at 37° C, media were replaced with PBS, and live cells imaged as described below.

For immunofluorescence, HepG2 cells were transfected with the pTarget-*TMEM135* construct. At 48 hours post-transfection, cells were fixed for 10 min at 37°C with 4% paraformaldehyde in culture media. Cells were washed twice with PBS, and permeabilized with 0.2% (*v:v*) Triton X-100 in PBS for 10 min at room temp. Cells were again washed in PBS, and incubated sequentially with Image-iT™ FX signal enhancer (Life Technologies) and 1% (*v:v*) normal goat serum in PBS for 30 min at room temperature. Antibodies against TMEM135 (Abcam catalog ab222237, rabbit polyclonal, 4 μg/ml) and ABCD3 (Abcam catalog ab211533, mouse monoclonal, 1 μg/ml) or COX4I1 (Abcam catalog ab33985, mouse monoclonal, 1 μg/ml) were diluted in PBS + 1% goat serum and incubated with cells overnight at 4°C. The next day cells were washed three times with PBS and incubated for two hours with DyLight® 488 donkey anti-rabbit IgG (ThermoFisher Scientific, 2 μg/ml) and Alexa Fluor®-594 goat anti-mouse IgG (Life Technologies, 4 μg/ml). Cells were washed and nuclei were stained with DAPI (600 nM). All imaging was performed on an EVOS fluorescent microscope (Life Technologies) equipped with a 60x objective (numerical aperture or NA = 0.75) and fluorescent light cubes for blue (peak excitation/emission or ex/em 357/447), green (ex/em 470/510), and red (ex/em 585/624) fluorescence.

### Gas chromatography (GC)

Lysates were prepared by sonicating frozen powdered liver in PBS as described earlier. Insoluble material was pelleted by centrifugation at 600 × g for 5 minutes. A BCA protein assay was performed on the cleared lysate. A one-step transesterification reaction was performed as described elsewhere [21]. Tridecanoic acid (13:0, Cayman Chemical) was used as an internal standard (10 μg/sample) because preliminary studies determined that it was not present in liver lysates. Authentic standards for the following fatty acids were obtained from Cayman Chemical: 12:0, 14:0, 16:0, 16:1, 18:0, 18:1 cis(n9), 18:2 cis(n6), 20:3 cis(n6), 20:4 (n6), 22:0, 22:6 (n3), 24:0, 24:1, and 26:0. Pooled mixtures containing increasing concentrations of these fatty acids were prepared and analyzed in parallel with liver samples. A total of 800 μg (protein content) of lysate from each sample underwent transesterification. Hexane was used to extract fatty acid methyl esters followed by evaporation under a gentle stream of nitrogen, and dried extracts were dissolved in 50 μl hexane.

GC was performed on an Agilent Technologies 6890N GC equipped with a Varian CP-Sil 88 column for fatty acid methyl ester analysis (100 m × 0.25 mm inner diameter × 0.2 μm film thickness) and a flame ionization detector. Inlet temperature was 250° C, and 1 μl of each standard or unknown was manually injected at a split ratio of 7.5:1. Helium was used as carrier gas at a 1.0 ml/minute constant flow. The oven was programmed for an initial temperature of 80° C with a 4° C/min ramp to 220° C, a 5-minute hold, then a 4° C/min ramp to 240° C followed by a 10-minute hold. The detector was set at 270°C using air (450 ml/minute) and hydrogen (40 ml/minute), and nitrogen was used as make-up gas (10 ml/minute constant flow). The sampling frequency was 20 Hz.

OpenLAB CDS ChemStation Edition software version C.01.06 (Agilent Technologies) was used to analyze GC data. Retention times of authentic standards were matched with corresponding peaks in unknown samples. For each fatty acid, the peak area for standards and unknowns was divided by the internal standard peak area. The normalized peak area from increasing concentrations of authentic standards was plotted, and the R^2^ for the resultant standard curves were > 0.995 in all cases. All fatty acids except 12:0 and 26:0 were identified in the unknowns, and fatty acid concentrations were determined by extrapolating the normalized peak area from the respective standard curve. Concentrations were expressed on an equal protein basis.

### Statistical analysis

Statistical analyses were performed using Stata Version 14 and all differences were considered significant at p< 0.05. For immortalized cell lines, the total sample size (n) corresponds to cells on a different passage number and/or experiments performed on different days. Time course data in immortalized cell lines were analyzed by mixed effects regression analysis to account for repeated measures. Treatments were coded 0 or 1, and hours post-treatment and treatment × time interaction were included as fixed effects in the regression model with experimental replicate as the random effect. Differences between treatments at each timepoint were determined from the treatment × time interaction coefficient. For all other comparisons involving more than 2 groups, mixed effects regression analysis was performed using treatment as the fixed effect and experimental replicate as the random effect to account for repeated measures. Pairwise differences between treatments were then determined using the Bonferroni multiple comparison method to control for the type 1 error rate. For comparisons involving only 2 groups, a paired t-test was performed.

For all mice data, the sample size refers to the number of mice. Regression analysis was used for mice outcomes except where indicated otherwise. Samples were coded 0 or 1 for fasting, TMEM135 knockdown fed mice, and TMEM135 knockdown fasted mice. The regression model utilized fasting, TMEM135 knockdown in the fed state, and TMEM135 knockdown in the fasted state, as predictors. The fasting coefficient indicated the difference due to fasting in control mice only, while the TMEM135 knockdown coefficients indicated the difference between control mice and TMEM135 knockdown mice within the fed state or fasted state. Differences were considered significant at p< 0.05, with p values ≥ 0.05 and < 0.10 considered trends. Only fed mice were used in proteomic analysis. Because the proteomic data does not follow a normal distribution due to some proteins not being detected in all samples, significant differences in proteomic data were determined from weighted spectra using the non-parametric Fisher’s Exact test (p < 0.05) contained within the Scaffold version 4.8.1 software. The list of proteins that were significantly reduced in knockdown livers was further trimmed by hand to exclude proteins that were not detected in all the samples from mice that received the control siRNA.

## Results

### LXR agonist effects on global gene expression in rhesus macaque luteal cells

When accounting for genes with multiple probe sets on the Affymetrix array, a total of 13 unique genes met the criteria for differential expression in rhesus macaque luteal cells, with 12 increased and 1 decreased by the LXR agonist T09 (see supplementary data for Excel spreadsheet of differentially expressed probe sets). The majority of differentially expressed genes were previously known LXR target genes including: *ABCA1* [5], *ABCG1* [6], *MYLIP* [7], *SREBF1* that encodes SREBP1c [8,9], *NR1H3* [10,11], *ACSL3* [22], *LPCAT3* [23], and *SCD* that is induced secondary to SREBP1c [24]. One differentially expressed gene that has not previously been linked to the LXRs was *TMEM135*, which was increased 2.55-fold by T09. A 4 kb region spanning from −3319 to +479 bp (relative to the translational start site) of the human *TMEM135* promoter was analyzed with MatInspector [25] for potential LXR binding sites. This analysis yielded three potential LXREs (Fig S1). Therefore, microarray analysis revealed *TMEM135* as a potentially novel direct LXR target gene.

### LXR agonist increases expression of TMEM135 in human hepatocyte and macrophage cell lines

Macrophage and hepatocyte cell lines were selected for further studies on LXR regulation of *TMEM135* transcription as these cell types play a pivotal role in LXR-mediated reverse cholesterol transport from peripheral tissues to the liver [1]. The LXR ligand T09 increased the mRNA expression of *TMEM135* in both HepG2 (Fig 1A) and Hep3B (Fig 1B) hepatocyte cell lines, with an approximately 2-fold maximum increase in HepG2 cells and a 3-fold increase in Hep3B cells. In monocyte-derived macrophages (THP-1), T09 also induced *TMEM135* mRNA expression up to 4-fold (Fig 1C). The protein synthesis inhibitor cycloheximide was used to determine whether the effect of T09 on *TMEM135* in THP-1 cells requires synthesis of new proteins. Cycloheximide did not significantly affect basal or T09-stimulated *TMEM135* mRNA expression (Fig 1D), indicating that T09 induces *TMEM135* via a direct transcriptional mechanism not involving the synthesis of intermediary proteins. Similar to mRNA, relative concentrations of TMEM135 protein were also significantly (p< 0.05) increased nearly 2-fold following a 48-hour T09 treatment in HepG2 cells (Fig 1E and 1F).

**Fig 1.**
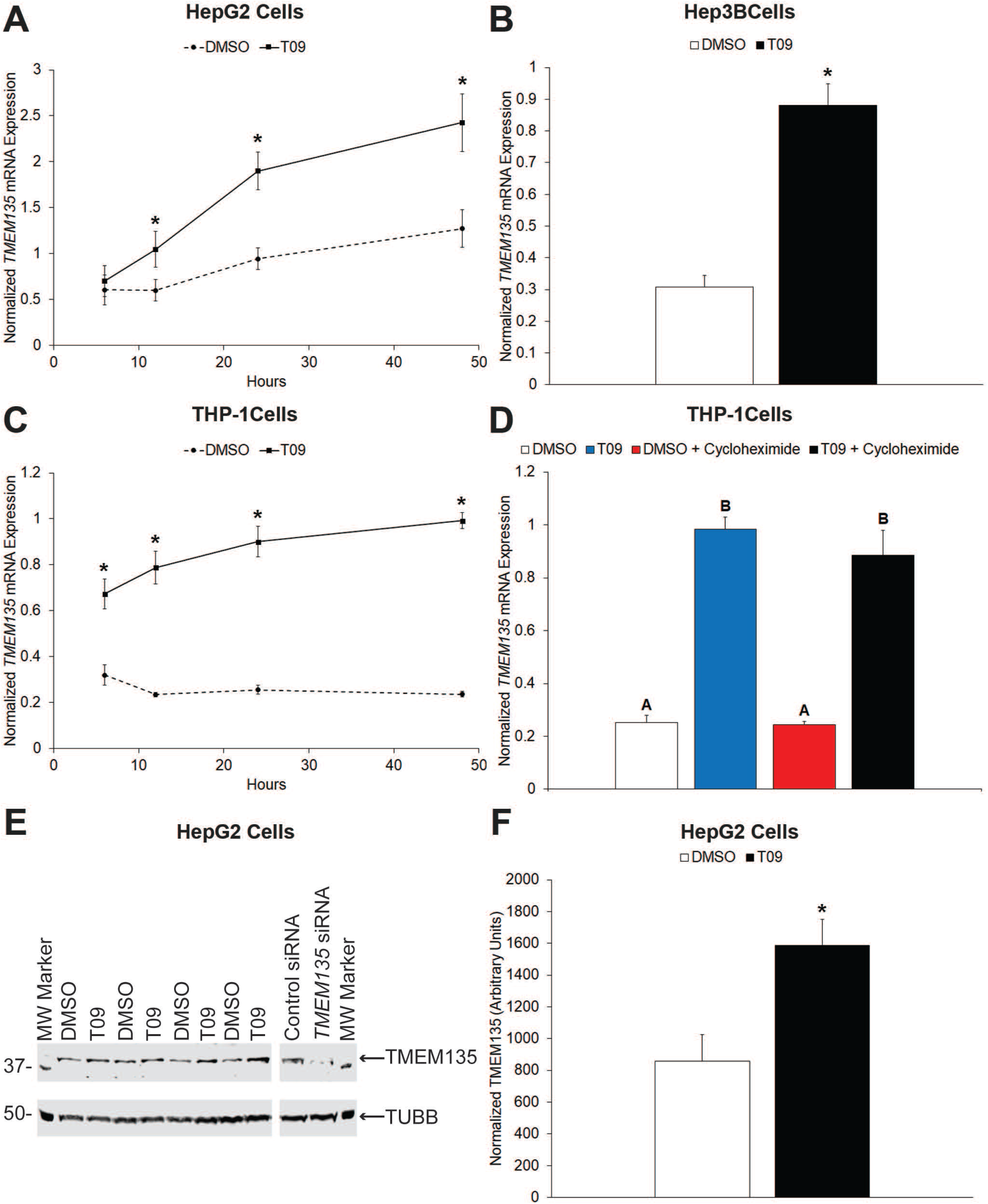
A synthetic LXR agonist induces TMEM135 expression in human hepatocyte (HepG2, Hep3B) and monocyte-derived macrophage (THP-1) cell lines via a direct transcriptional mechanism. Panel A contains the results from treatment of HepG2 cells over time in the presence or absence of the synthetic LXR agonist T09; while panel B is the effect of a 24-hour T09 treatment on *TMEM135* mRNA expression in Hep3B cells (n = 4). Panel C is a time course of T09 treatment in THP-1 cells (n = 4). For panels A-C, asterisks denote significant (p< 0.05) difference between DMSO and T09 at the indicated timepoint. Panel D is the effect of the protein synthesis inhibitor cycloheximide on *TMEM135* mRNA expression in THP-1 cells. Cells were treated for 24 hours in a 2 × 2 factorial with T09 and cycloheximide (n = 5). Columns with different letters are significantly different (p< 0.05). Panel E is Western blot analysis of TMEM135 in HepG2 cells treated with DMSO or T09 for 48 hours (n = 4). The approximate molecular weights of the molecular weight (MW) marker are indicated, as well as the TMEM135 and TUBB (housekeeping control) bands. As a validation of TMEM135 band identity, lysates from HepG2 cells transfected with a control or *TMEM135* siRNA were also run. Panel F is results of densitometry analysis of the blot shown in Panel E, asterisk denotes significant difference (p< 0.05). In panels A-D, the normalized value is the ratio of *TMEM135* to the housekeeping gene *MRPS10*. In panel F, TMEM135 is normalized to TUBB. For all panels, error bars indicate ± one standard error of the mean (SEM).

### Identification of the LXRE that mediates LXR agonist induction of *TMEM135*

The potential LXREs identified by MatInspector were arbitrarily designated LXRE1, LXRE2 and LXRE3 in order from most distal to most proximal to the translation start site (Fig S1). Electrophoretic mobility shift assays (EMSA) were used to determine if LXR/RXR heterodimers bind these LXREs. As shown in Fig 2A, NR1H3 and RXRA nuclear receptors together, but not individually, caused a shift in mobility of the fluorescent LXRE1 probe indicating that NR1H3/RXRA heterodimers bind the LXRE1 sequence in the *TMEM135* promoter. A 200-fold molar excess of unlabeled LXRE1, LXRE2, and LXRE3 oligonucleotides eliminated the appearance of the shifted fluorescent band while the same molar excess of mutant LXRE1, LXRE2, and LXRE3 oligonucleotides (Fig S1B) did not prevent NR1H3/RXRA binding to the fluorescent probe; which validated the specificity of NR1H3/RXRA heterodimer binding to all 3 LXREs in the *TMEM135* promoter (Fig 2A). Similar results were obtained for NR1H2/RXRA heterodimers (Fig 2B), indicating that both LXR isoforms specifically bind to all 3 LXRE sites in the *TMEM135* promoter. The LXRE1 and LXRE2 sites have identical hexanucleotide half-sites, whereas the LXRE3 site is unique (Fig S1B). To indicate if these sequence variations may cause differences in relative binding affinity, a fluorescent LXRE3 probe was incubated with NR1H3/RXRA heterodimers and increasing concentrations (from 0.25 to 10-fold molar excess) of either LXRE1 or LXRE3 unlabeled competitor DNA. Oligonucleotides with the LXRE3 sequence more effectively inhibited binding to the fluorescent probe than oligonucleotides with the LXRE1 sequence (Fig 2C). As a complementary approach, because the LXRE1 and LXRE3 sites were fluorescently labelled with spectrally distinct fluorophores, an EMSA was performed where a mixture of fluorescent LXRE1 and LXRE3 probes were incubated with increasing concentrations of NR1H3/RXRA heterodimers. This approach resulted in a higher percentage of LXRE3 than LXRE1 bound across a range of NR1H3/RXRA heterodimer concentrations (Fig 2D). Collectively, this indicates that LXRE3 has a higher binding affinity for NR1H3/RXR heterodimers than LXRE1.

**Fig 2.**
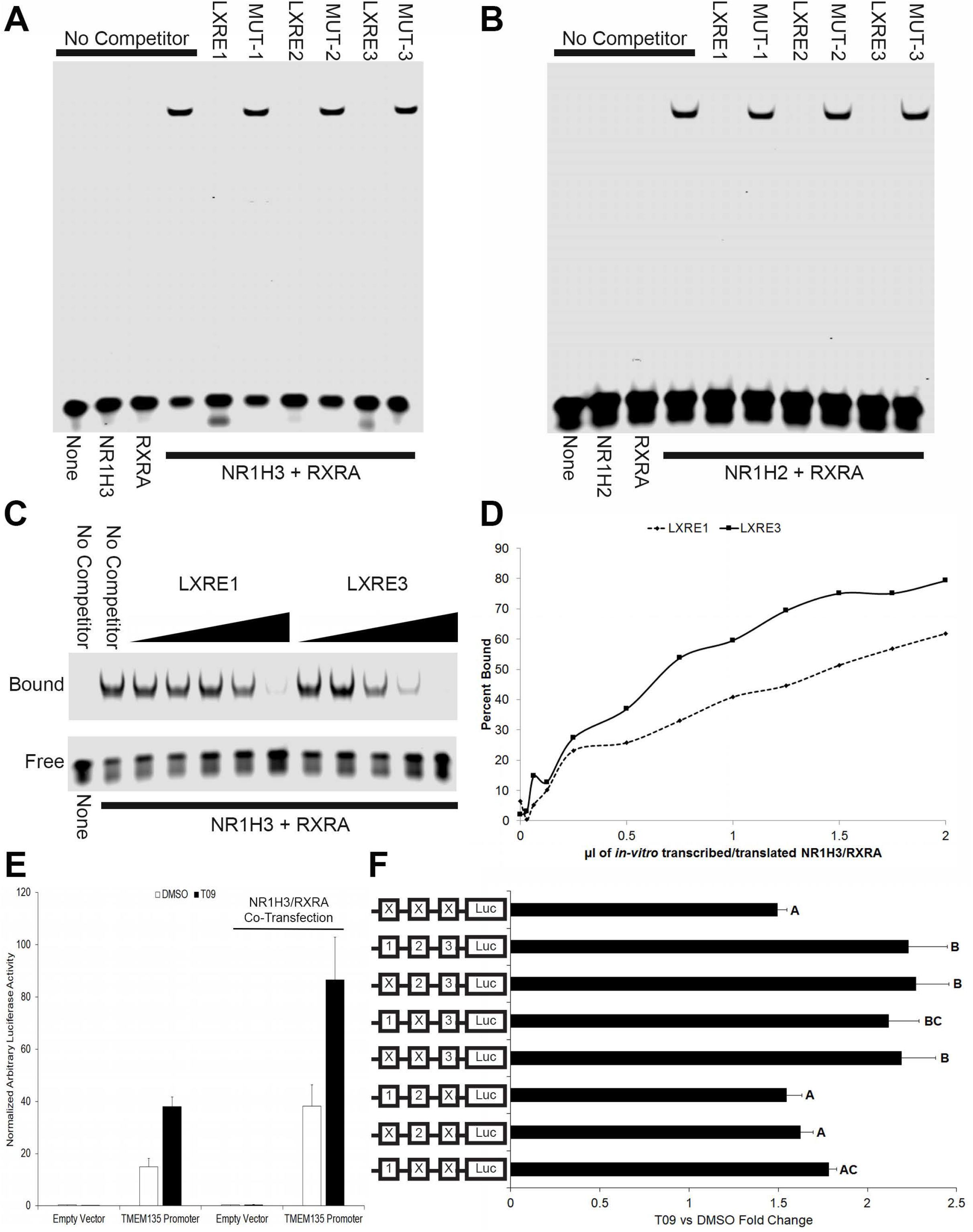
The LXRs bind all three potential LXREs in the human *TMEM135* promoter with LXRE3 mediating LXR-induced *TMEM135* transcription. Panel A is an EMSA using LXRE1 as the fluorescent probe. The nuclear receptor(s) used in the reaction are indicated beneath the image, with the unlabeled competitor DNA (200-fold molar excess) shown above the image. Panel B substitutes NR1H2 for NR1H3 in the binding reactions. Panel C utilizes fluorescent LXRE3 as the probe. The image has been cropped to only show the bound and free fluorescent LXRE3 probe. The nuclear receptor(s) used in the reaction are indicated beneath the image. Increasing amounts (0.25, 0.5, 1, 2, and 10-fold molar excess per competitor) of unlabeled LXRE1 or LXRE3 were included in some reactions as indicated above the images. In Panel D, an EMSA was performed using a mixture of LXRE1 and LXRE3 probes that were labeled with spectrally-distinct fluorescent dyes, as well as increasing quantities of LXR/RXR proteins. The percent of each probe bound in each reaction was determined by densitometry and plotted. Panel E displays luciferase activity (arbitrary units normalized to β-galactosidase) derived from cells transfected with the empty vector or the *TMEM135* promoter-containing construct in the presence and absence of T09. The effect of increased expression of NR1H3 and RXRA is also shown (n = 4). Panel F contains the fold-increase in luciferase activity induced by T09 from the wild type *TMEM135* promoter, as well as *TMEM135* promoters containing all possible combinations of mutant LXRE sites (n = 4). All transfections in Panel F included *NR1H3* and *RXRA* co-transfection to increase basal expression of these nuclear receptors. An X indicates point mutations were introduced into the corresponding LXRE (see Fig S1), Luc = luciferase. Error bars indicate one SEM, columns without common letters are significantly (p< 0.05) different.

Luciferase reporter assays were used to study transcription initiation from the *TMEM135* promoter. Transfection of HepG2 cells with wild type *TMEM135* promoter/pGL 4.17 resulted in a large increase in luciferase expression that was responsive to T09 as compared to the empty pGL 4.17 vector (Fig 2E). Furthermore, luciferase activity was further amplified by co-transfection of plasmids that constitutively express NR1H3 and RXRA (Fig 2E). Next, we determined the requirement of each individual LXRE to LXR agonist-induced transcription from the *TMEM135* promoter. Promoters containing all combinations of wild type and mutant LXREs (Fig S1B) were generated, and T09-induced luciferase activity was determined. LXR-agonist induced luciferase activity was significantly (p< 0.05) higher in cells transfected with the wild type *TMEM135* promoter as compared to the *TMEM135* promoter with all 3 LXREs mutated (Fig 2F). Furthermore, constructs containing the wild type LXRE3 site and mutated LXRE1 and/or LXRE2 sites were not significantly different from the wild type promoter, while all constructs that contained a mutated LXRE3 were not significantly different from the promoter that had all three LXREs mutated (Fig 2F). This indicates that even though all 3 LXREs bind LXR/RXR heterodimers, the LXRE3 site alone mediates LXR agonist-induced transcription from the *TMEM135* promoter.

We next determined whether *Tmem135* mRNA expression was induced by T09 in mice. The murine LXRE3 site showed relatively little homology with the human sequence as a total of 3 nucleotides within both hexanucleotide half-sites were unique in mice (Fig S2A). In mouse hepatocyte (BNL 1NG A.2) and macrophage (RAW 264.7) cell lines, T09 did not significantly induce *Tmem135* mRNA expression (Fig S2B) whereas it did increase the known LXR target gene *Abca1* (Fig S2C). These data indicate that *Tmem135* is not an LXR target gene in mice.

### *TMEM135* is a direct target gene of the LXRs

To directly determine whether the LXRs are needed for T09-induced *TMEM135* transcription, siRNA-mediated knockdown of the LXRs was performed in HepG2 cells. The *NR1H2* siRNA caused an approximately 75% decrease (p< 0.05) in *NR1H2* mRNA expression compared to the control and *NR1H3* siRNA groups (Fig 3A). The *NR1H3* siRNA resulted in an approximately 70-80% decrease (p< 0.05) in *NR1H3* mRNA expression compared to the control siRNA depending on the presence or absence of T09 (Fig 3B). The *NR1H2* siRNA by itself also caused a significant reduction in T09-stimulated *NR1H3* mRNA expression and tended to slightly improve *NR1H3* knockdown when co-transfected with the *NR1H3* siRNA (Fig 3B). Decreased *NR1H3* mRNA expression is expected to occur following NR1H2 knockdown because *NR1H3* itself is an LXR target gene [6,11]. Consistent with *NR1H3* being an LXR target gene, T09 induced a significant increase in *NR1H3* in the control siRNA group, while T09 was not as effective at inducing *NR1H3* in the *NR1H2* siRNA group (Fig 3B).

**Fig 3.**
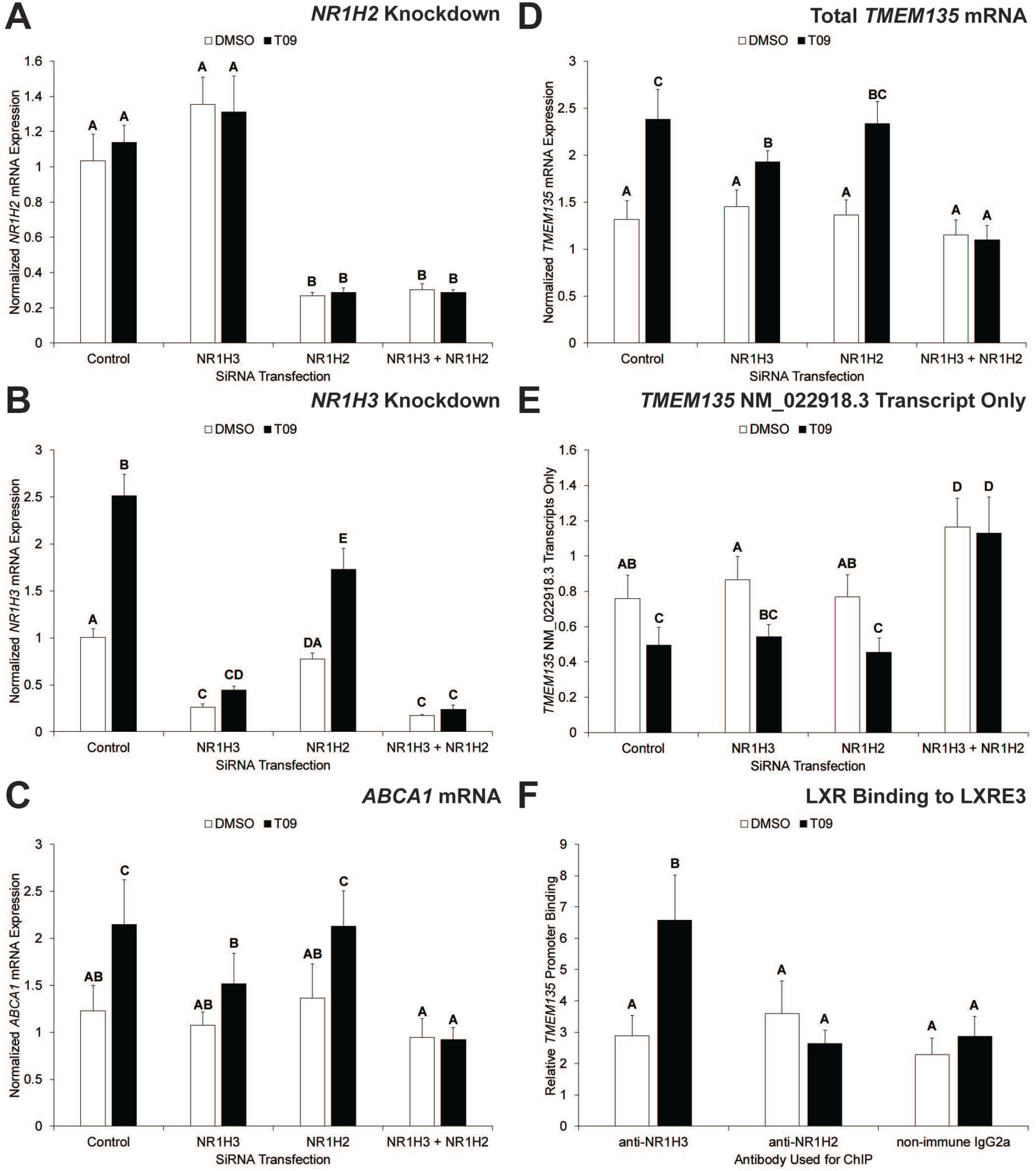
The LXRs are obligatory for LXR agonist-induced *TMEM135* mRNA expression, and cause transcription of *TMEM135* with an abbreviated 5’ UTR. Panels A, B, C, and D show the effects of siRNA transfection in HepG2 cells on T09-induced mRNA expression of the LXR isoforms *NR1H2* and *NR1H3*, *ABCA1*, and *TMEM135*; respectively. Panel E shows the effect of LXR knockdown in HepG2 cells on T09-induced expression of *TMEM135* mRNA transcripts containing a longer 5’ UTR (based on NM_022918.3). Primers and probe used for QPCR in panel E amplify a 76 bp region that encompasses the LXR binding site in genomic DNA, which is present in the NM_022918.3 transcript but absent from the NM_022918.4 transcript. Panel F is results from ChIP analysis. For panels A-E, data are normalized to *MRPS10*. For panel F, data are normalized to a non-specific locus. For all panels, error bars indicate one SEM, columns without common letters are significantly (p< 0.05) different (n = 4).

The known LXR target gene *ABCA1* was significantly increased by T09 in the control siRNA group, and the effect of T09 was completely abolished by *NR1H2/NR1H3* siRNA co-transfection (Fig 3C). Also, *NR1H3* knockdown alone significantly inhibited the T09-induced increase in *ABCA1* (Fig 3C). Results for *TMEM135* were very similar to *ABCA1* with T09 causing a significant increase in the control siRNA group, and the effect of T09 was completely blocked by *NR1H2/NR1H3* siRNA co-transfection (Fig 3D). Furthermore, *NR1H3* knockdown itself significantly inhibited the T09-induced increase in *TMEM135* as compared to the control siRNA (Fig 3D).

When these studies were originally performed the putative transcription and translation start sites of human *TMEM135* were based on NCBI accession number NM_022918.3 (Fig S1), which indicated that LXRE3 was in the 5’ untranslated region (5’ UTR). To determine whether LXR-mediated transcription of *TMEM135* resulted in an mRNA with a truncated 5’ UTR, primers and a probe were designed to amplify and detect a 76 base pair region encompassing the LXRE3 site. There was a significant (p< 0.05) reduction in mRNA expression of transcripts containing the LXRE3 site (NM_022918.3) in the control siRNA group, while this effect was blocked by *NR1H2/NR1H3* siRNA co-transfection (Fig 3E). Furthermore, *NR1H2/NR1H3* co-knockdown resulted in a significant (p< 0.05) increase in mRNA expression of transcripts containing the LXRE3 site compared to all other groups (Fig 3E). ChIP was used to determine if the reduction in transcripts containing the LXRE3 site is associated with increased NR1H2 and/or NR1H3 binding to the LXRE3 site. Monoclonal antibodies specific for each LXR isoform were used for ChIP, and QPCR of the LXRE3 region on chromatin purified via ChIP indicated that T09 treatment caused a significant increase in binding of NR1H3 to LXRE3 in the *TMEM135* gene (Fig 3F). On November 23, 2018 NCBI accession number NM_022918.4 replaced NM_022918.3, which identified a transcription start site downstream of LXRE3. This indicates that LXR binding to LXRE3 shifts mRNA expression of *TMEM135* from the NM_022918.3 transcriptional start site to the NM_022918.4 transcriptional start site. It should be noted that both transcripts have the same translation start site, but the truncated 5’ UTR in NM_022918.4 may alter overall TMEM135 protein expression (e.g., by altering post-transcriptional events such as miRNA binding), which should be explored in future studies.

### TMEM135 mediates fatty acid metabolism and proliferation in HepG2 cells

To begin unraveling the biologic function of TMEM135, a series of knockdown experiments were performed. Transfection of HepG2 cells with siRNA against *TMEM135* decreased its mRNA expression by 70-90%. As TMEM135 has previously been implicated in fat metabolism [26], and the LXRs are known to induce lipogenesis [1], we first determined its effect on triglycerides and mRNA expression of fatty acid oxidation and lipogenesis-associated genes. Knockdown of TMEM135 significantly increased basal triglyceride accumulation in HepG2 cells (Fig 4A). Furthermore, T09 itself significantly increased intracellular triglyceride concentrations, while TMEM135 knockdown further increased triglyceride accumulation in the presence of T09 (Fig 4A). Neutral lipid staining of HepG2 cells appeared consistent with the detected changes in intracellular triglycerides (Fig 4B). An increase in triglyceride accumulation could result from a decrease in fatty acid oxidation and/or an increase in lipogenesis. There were no significant effects of TMEM135 knockdown on mRNA expression of the key regulator of fatty acid β-oxidation, peroxisome proliferator activated receptor alpha (*PPARA*) or the PPARA target gene carnitine palmitoyltransferase 1A (*CPT1A*) [27] (Fig 4C). It is known that the LXRs induce lipogenesis via induction of SREBP1c (*SREBF1* gene) [8,9]. As expected, T09 caused a significant increase in *SREBF1* and the SREBP1c/LXR target gene fatty acid synthase (*FASN*) (Fig 4C), consistent with the T09-induced increase in triglyceride accumulation (Fig 4A). However, TMEM135 knockdown significantly suppressed both basal and T09-induced *SREBF1* mRNA expression (Fig 4C). This indicates that the increase in triglyceride accumulation in HepG2 cells occurred despite a reduction in lipogenic gene expression. Thus, an inhibition of fatty acid oxidation seems a more likely explanation for the increase in triglycerides.

**Fig 4.**
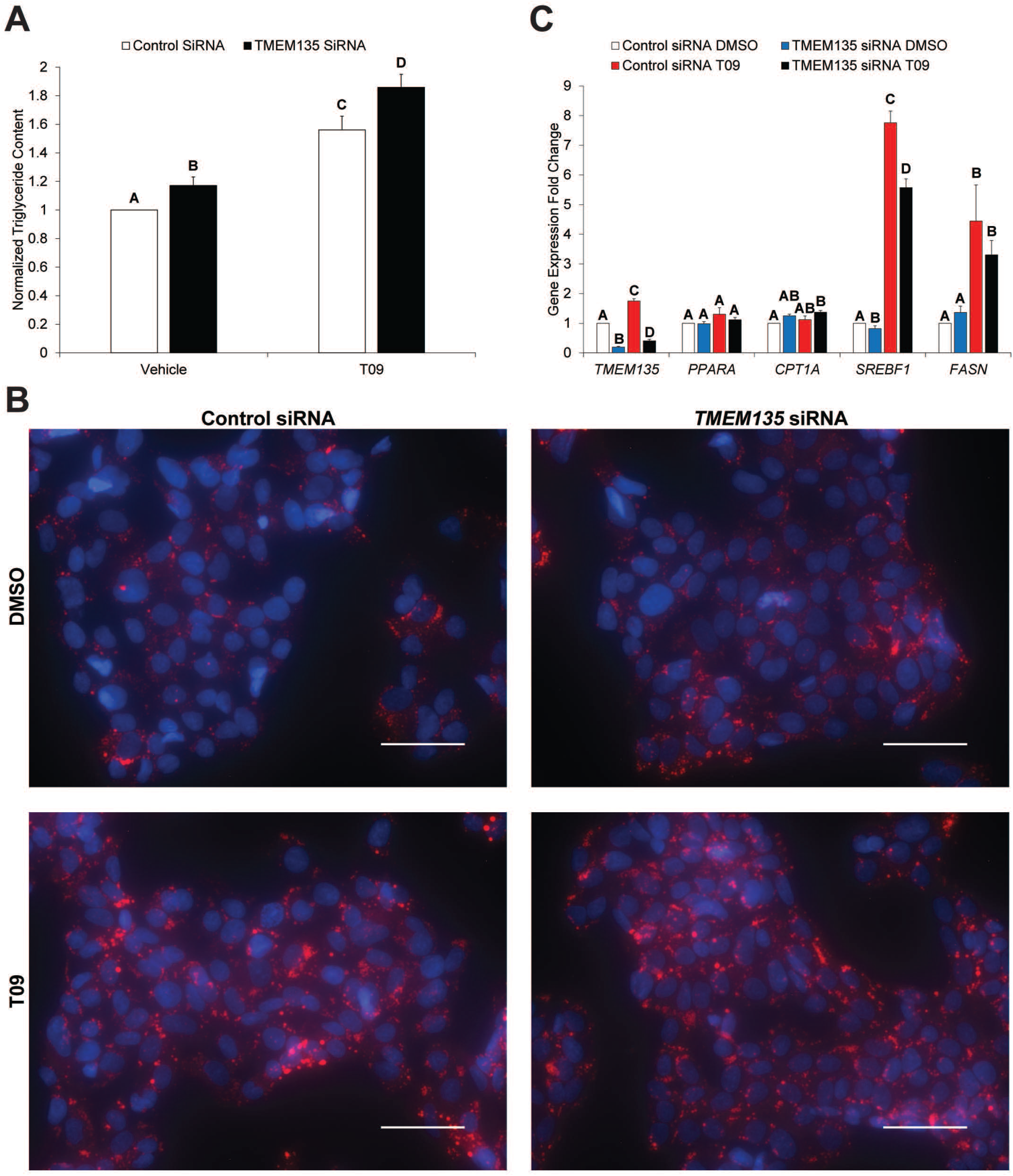
TMEM135 regulates fatty acid metabolism in HepG2 cells. Panel A displays the effect of TMEM135 knockdown in the presence and absence of the lipogenic LXR agonist T09 on intracellular triglyceride accumulation in HepG2 cells. Triglyceride concentrations were normalized to protein concentrations, and for presentation purposes the data are plotted as a fold-change relative to the control siRNA + vehicle group. Panel B is neutral lipid staining (red) of HepG2 cells with the same transfections and treatments as in panel A. Nuclei are blue, and the scale bar represents 50 μm. Panel C is mRNA expression in HepG2 cells for *TMEM135*, genes involved in fatty acid oxidation (*PPARA*, *CPT1A*), and genes involved in lipogenesis (*SREBF1*, *FASN*). All genes were normalized to *MRPS10* and plotted as a fold-change relative to the control siRNA + vehicle group. For panels A and C, columns without a common letter are significantly (p< 0.05) different (n = 4).

During these experiments it appeared that TMEM135 knockdown also inhibited replication of HepG2 cells. Because enhanced β-oxidation is a hallmark of hepatocellular carcinoma (HCC) [28] and our previous experiments indicated a key role for TMEM135 in fatty acid metabolism, we determined whether TMEM135 regulated proliferation of HepG2 cells. Knockdown of *TMEM135* significantly (p< 0.05) reduced viable HepG2 cell numbers at 48 and 72-hours post-transfection as compared to cells transfected with the control siRNA (Fig S3A). Cell cycle analysis indicated that *TMEM135* knockdown significantly (p< 0.05) increased the percentage of HepG2 cells in the G0/G1 stage, with a corresponding significant reduction in the percentage of cells in the S phase (Fig S3B). The increase in G0/G1 arrest was associated with alterations in mRNA expression of tumor suppressor and cell cycle genes. There were significant increases in cyclin dependent kinase inhibitor 2A (*CDKN2A*) and tumor protein p53 (*TP53*) (Fig S3C), which restrict passage through the G1/S checkpoint [29]. There were also significant increases in cyclin dependent kinase 2 (*CDK2*) and cyclin E1 (*CCNE1*) (Fig S3C), which increase prior to passage through the G1/S checkpoint, but no change in cyclin A2 (*CCNA2*) (Fig S3C) which is increased in the S phase [30]. Collectively, these data indicate that TMEM135 knockdown reduces HepG2 proliferation by restricting passage through the G1/S checkpoint. Further supporting an impairment in fatty acid β-oxidation that could reduce proliferation, TMEM135 knockdown significantly reduced ATP concentrations when HepG2 cells were incubated in glucose-free medium (Fig S3D).

### Liver-selective TMEM135 knockdown reduces peroxisomal β-oxidation

To determine the physiologic function of TMEM135, a siRNA knockdown experiment was performed in male C57BL/6 mice. Mice received either a non-targeting control or *Tmem135* siRNA and were sacrificed 4 days later in either the *ad libitum* fed state or after a 12-hour fast. Because we determined that *Tmem135* is not an LXR target gene in mice (Fig S2), we used fasting to acutely induce hepatic fat accumulation. The *Tmem135* siRNA caused an approximately 60% mRNA knockdown in the liver while no knockdown was observed in other tissues including skeletal muscle, adipose, and heart (Fig 5A). Knockdown of TMEM135 in the liver was further confirmed by Western blot analysis (Fig 5B). Fed mice injected with the *Tmem135* siRNA gained significantly less weight during the 4-day treatment, while the loss in weight from fasting was similar for both siRNAs (Fig 5C). There was no significant effect of the *Tmem135* siRNA on basal or fasting-induced hepatic triglyceride, ATP, or glycogen concentrations (Fig S4A-C). Also, there were no significant effects of TMEM135 on serum lipids (total cholesterol, HDL cholesterol, triglycerides), although there was a trend for fasting to reduce serum triglycerides in control siRNA mice that was not observed in TMEM135 knockdown mice (Fig S5A). There were no significant effects of TMEM135 knockdown on serum NEFA, glucose, insulin, or β-hydroxybutyrate concentrations (Fig S5B-E). The mRNA expression of key genes involved in fatty acid β-oxidation were determined in the liver. As expected, fasting significantly induced mRNA expression of *Ppara*, *Cpt1a*, acyl-CoA dehydrogenase medium chain (*Acadm*), uncoupling protein 2 (*Ucp2*), and sirtuin 3 (*Sirt3*) (Fig 5D). Similarly, knockdown of TMEM135 significantly increased *Acadm* and *Sirt3* in fed animals, while *Ucp2* was significantly reduced by TMEM135 knockdown in fasted animals (Fig 5D). As expected with increased β-oxidative flux, fasting significantly increased NADH in animals receiving the control siRNA, while TMEM135 knockdown significantly reduced NADH (Fig 5E) indicating an impairment in β-oxidation in fasted mice. Reduced hepatic NADH during fasting could be due to reduced β-oxidation in peroxisomes and/or mitochondria. However, during fasting hepatic β-hydroxybutyrate concentrations in TMEM135 knockdown mice (Fig 5F) tended (p = .098) to increase compared with controls. Because ketogenesis occurs exclusively in mitochondria and is intricately linked with fatty acid oxidation [31], this indicates that mitochondrial fatty acid β-oxidation was not impaired. Collectively, NADH and ketone data in the fasted state are consistent with an impairment in peroxisomal β-oxidation.

**Fig 5.**
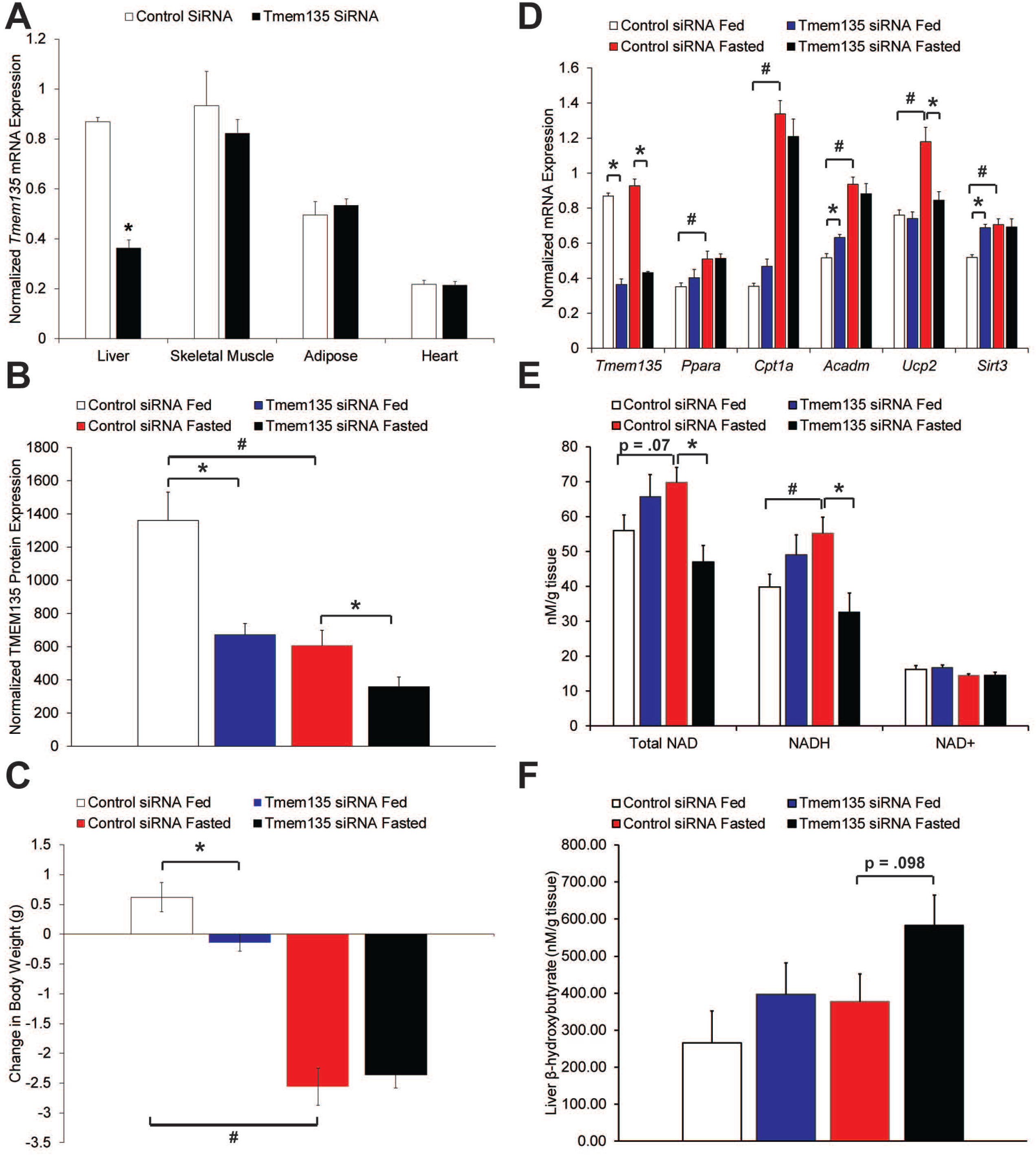
Liver-specific knockdown of TMEM135 inhibits peroxisomal β-oxidation. Panel A is mRNA expression of *Tmem135* normalized to *Mrps10* in the listed tissues isolated from fed mice (n = 5). Asterisk denotes significant difference due to *Tmem135* siRNA. Panel B displays TMEM135 protein expression normalized to TUBB in livers. Panel C displays the change in body weight of animals from the time of siRNA injection until sacrifice 4 days later. Panel D displays hepatic mRNA expression of *Tmem135* and genes involved in fatty acid oxidation normalized to *Mrps10*, panel E displays hepatic NAD concentrations, and Panel F is hepatic β-hydroxybutyrate concentrations. For panels B-F, asterisks denote significant difference (p< 0.05) due to *Tmem135* siRNA within feeding status, and # indicates significant difference due to fasting in animals receiving control siRNA (n = 5 per siRNA and nutritional status).

### TMEM135 knockdown reduces matrix enzyme concentrations within peroxisomes

TMEM135 is a peroxisomal protein with homology to the Tim17 family that mediate translocation of proteins across mitochondrial membranes [32]. In addition to peroxisomes, TMEM135 has been reported to be localized to mitochondria [26,33]. Therefore, we hypothesized that TMEM135 mediates enzyme concentrations within peroxisomes and/or mitochondria. A mitochondria/peroxisome-enriched fraction was prepared from frozen mouse livers. This fraction was validated to contain both mitochondria and peroxisomes as indicated by presence of the mitochondria marker cytochrome c oxidase subunit 4I1 (COX4I1) and the peroxisome marker ATP binding cassette subfamily D member 3 (ABCD3, also known as PMP70), while COX4I1 and ABCD3 were not detected in the cytosolic fraction (Fig 6A). Furthermore, the cytosolic protein tubulin beta class I (TUBB) segregated to the cytosolic fraction and was not detected in the mitochondria/peroxisome-enriched fraction. Proteomic analysis of a subset of fed mice was used to provide an unbiased estimate of differential protein abundance in the mitochondria/peroxisome-enriched fractions. A total of 23 proteins were significantly less abundant (Fisher’s Exact Test, p< 0.05) in the mitochondria/peroxisome-enriched fraction from TMEM135 knockdown livers (Table 1). Interestingly, these 23 proteins included nearly all the matrix proteins known to be necessary for peroxisomal bile acid synthesis and β-oxidation of fatty acids, branched chain fatty acids, and dicarboxylic acids [34,35]. The peroxisomal membrane protein ABCD3 was used as a marker of total peroxisome content. Concentrations of ABCD3 determined via proteomic and Western blot analysis demonstrated a tendency for reduced peroxisome content following TMEM135 knockdown, but the differences were not statistically significant (Fig 6B).

**Fig 6.**
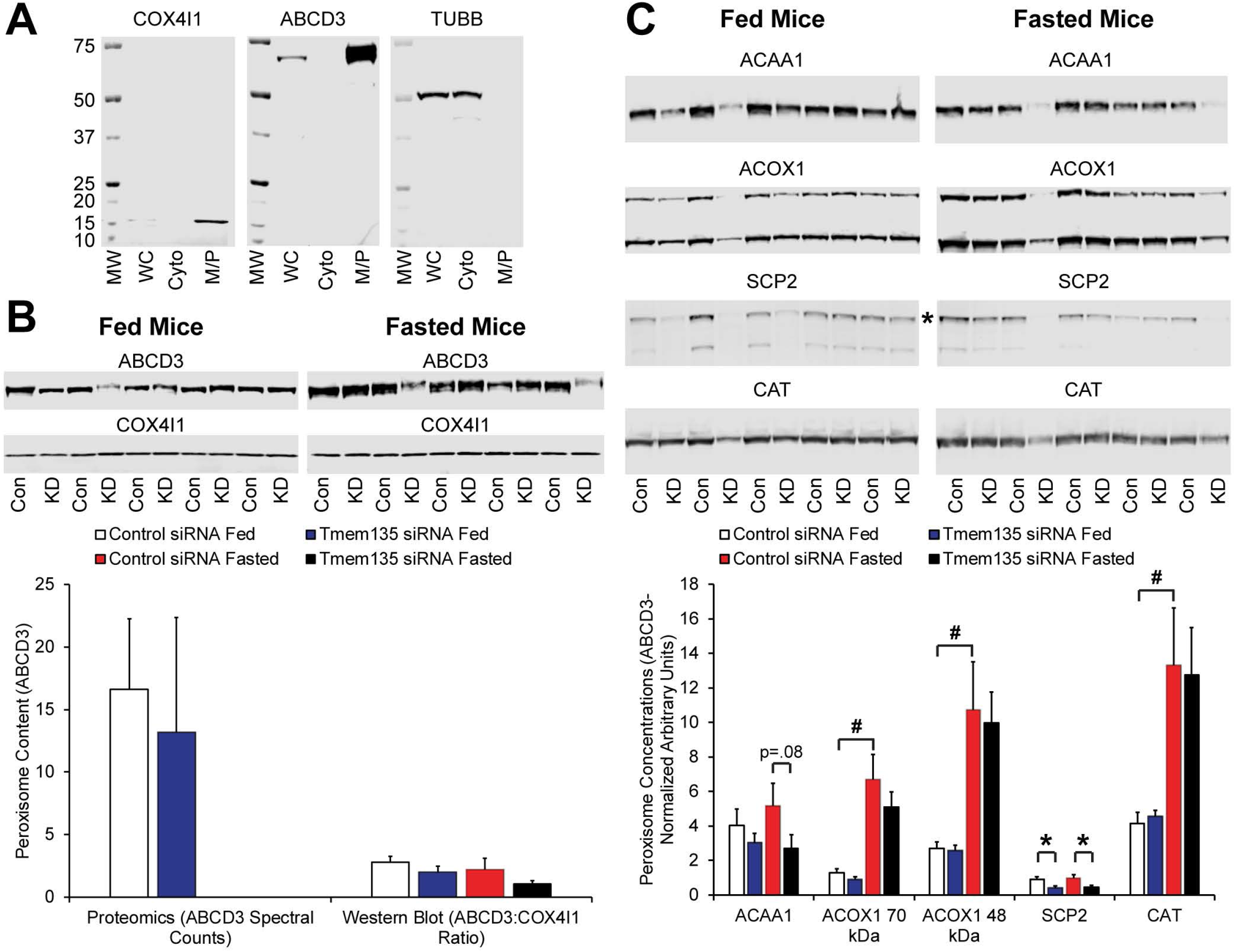
TMEM135 regulates peroxisomal matrix enzyme concentrations. Panel A displays the relative purity of the mitochondria/peroxisome-enriched fraction used for proteomic analysis as determined by Western blot analysis of controls for mitochondria (COX4I1), peroxisome (ABCD3), and cytosolic (TUBB) proteins. For each image, the lanes from left to right are the molecular weight marker (MW), whole cell lysate (WC), cytosolic fraction (Cyto), and the mitochondria/peroxisome-enriched fraction (M/P) from a control siRNA treated, fasted mouse. Panel B displays relative peroxisome content in the mitochondria/peroxisome-enriched fractions as determined by proteomic and Western blot analysis of ABCD3. For proteomics, only fed mice were analyzed (n = 3). For Western blot, the ABCD3:COX4I1 ratio in the mitochondria/peroxisome-enriched fraction is shown (n = 5 per siRNA and nutritional status). Panel C contains images from Western blot analysis of mitochondria/peroxisome-enriched fractions in livers of fed and fasted mice. The siRNA treatment is indicated beneath the lanes, Con = control siRNA, KD = Knockdown, *Tmem135* siRNA. ACOX1 migrates in 2 separate bands in Western blot, with the smaller band being proteolytically processed ACOX1 [36]. Furthermore, SCP2 (also known as SCPx) is a 58 kDa protein that contains a 45 kDa 3-ketoacyl CoA thiolase domain and a 13 kDa sterol-carrier domain [37], which are separated by proteolytic cleavage [36]. The antibody used in the current study recognizes the full-length 58 kDa protein (indicated by asterisk) and the 45 kDa 3-ketoacyl CoA thiolase domain (lower band), but not the 13 kDa sterol-carrier domain. The 58 kDa band was used for densitometry. Results of densitometry analysis is shown beneath the blots. Asterisks denote significant difference (p< 0.05) due to *Tmem135* siRNA within feeding status, and # indicates significant difference due to fasting in animals receiving control siRNA (n = 5 per siRNA and nutritional status).

**Table 1.**
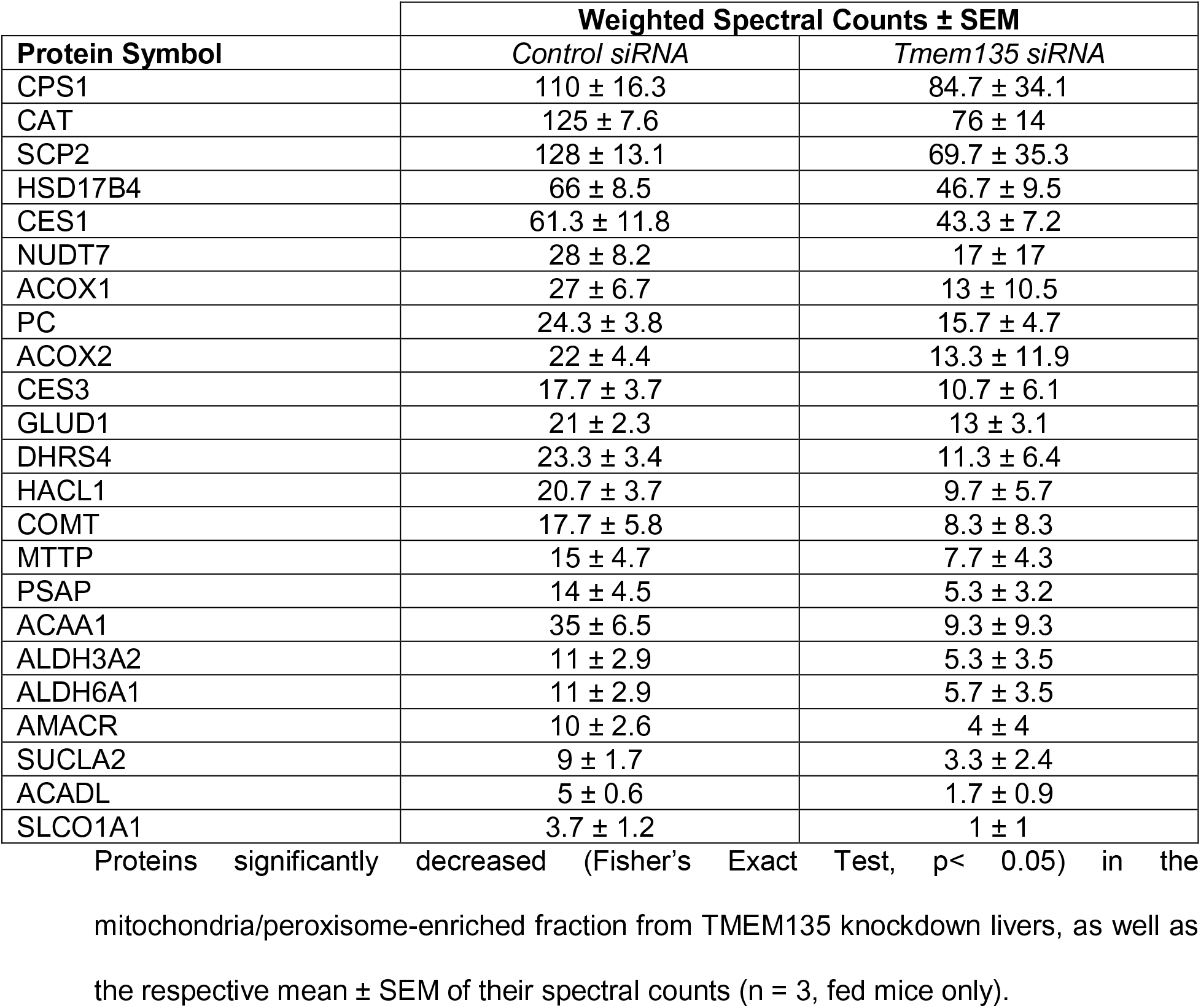
Proteomic analysis of mitochondria/peroxisome-enriched fraction.

| **Protein Symbol** | **Weighted Spectral Counts ± SEM** |  |
| --- | --- | --- |
|  | *Control siRNA* | *Tmem135 siRNA* |
| CPS1 | 110 ± 16.3 | 84.7 ± 34.1 |
| CAT | 125 ± 7.6 | 76 ± 14 |
| SCP2 | 128 ± 13.1 | 69.7 ± 35.3 |
| HSD17B4 | 66 ± 8.5 | 46.7 ± 9.5 |
| CES1 | 61.3 ± 11.8 | 43.3 ± 7.2 |
| NUDT7 | 28 ± 8.2 | 17 ± 17 |
| ACOX1 | 27 ± 6.7 | 13 ± 10.5 |
| PC | 24.3 ± 3.8 | 15.7 ± 4.7 |
| ACOX2 | 22 ± 4.4 | 13.3 ± 11.9 |
| CES3 | 17.7 ± 3.7 | 10.7 ± 6.1 |
| GLUD1 | 21 ± 2.3 | 13 ± 3.1 |
| DHRS4 | 23.3 ± 3.4 | 11.3 ± 6.4 |
| HACL1 | 20.7 ± 3.7 | 9.7 ± 5.7 |
| COMT | 17.7 ± 5.8 | 8.3 ± 8.3 |
| MTTP | 15 ± 4.7 | 7.7 ± 4.3 |
| PSAP | 14 ± 4.5 | 5.3 ± 3.2 |
| ACAA1 | 35 ± 6.5 | 9.3 ± 9.3 |
| ALDH3A2 | 11 ± 2.9 | 5.3 ± 3.5 |
| ALDH6A1 | 11 ± 2.9 | 5.7 ± 3.5 |
| AMACR | 10 ± 2.6 | 4 ± 4 |
| SUCLA2 | 9 ± 1.7 | 3.3 ± 2.4 |
| ACADL | 5 ± 0.6 | 1.7 ± 0.9 |
| SLCO1A1 | 3.7 ± 1.2 | 1 ± 1 |

Proteins significantly decreased (Fisher's Exact Test, $p < 0.05$ ) in the mitochondria/peroxisome-enriched fraction from TMEM135 knockdown livers, as well as the respective mean ± SEM of their spectral counts ( $n = 3$ , fed mice only).

Four peroxisomal matrix proteins were selected for further analysis via Western blot: acetyl-CoA acyltransferase 1 (ACAA1), acyl-CoA oxidase 1 (ACOX1), sterol carrier protein 2 (SCP2, also known as SCPx), and catalase (CAT). Because their was some variability in the peroxisome content of the mitochondria/peroxisome-enriched fraction (Fig 6B), the signal for each protein was normalized to ABCD3 to adjust for differences in total peroxisome content. There were no significant differences between siRNAs for peroxisomal concentrations of ACOX1 and CAT in either fed or fasted animals (Fig 6C), indicating their decrease in the proteomics dataset was at least partially due to a reduced peroxisome content. However, peroxisomal SCP2 concentrations were significantly lower in TMEM135 knockdown mice in both the fed and fasted state, while ACAA1 concentrations tended (p = .08) to be lower in TMEM135 knockdown mice in the fasted state (Fig 6C). This indicates that TMEM135 increases the concentrations of SCP2 and ACAA1 within intact peroxisomes in the livers of fed and/or fasted mice independent of changes in peroxisome number, and thus TMEM135 may regulate intraperoxisomal concentrations of matrix enzymes.

### Localization of TMEM135

Liver NADH and ketone concentrations (Fig 5), as well as proteomic data (Table 1), indicated that TMEM135 is functioning in peroxisomes. This contradicts two previous studies that reported TMEM135 localization to mitochondria [26,33]. Therefore, we sought to determine if TMEM135 is localized to peroxisomes and/or mitochondria. A TMEM135-EGFP fusion plasmid was co-transfected in HepG2 cells with a vector expressing mCherry with a PTS1 peroxisome targeting sequence, or a vector expressing mCherry with a mitochondrial localization sequence. The TMEM135-EGFP fusion protein appeared as punctate spots in the cytoplasm which co-localized with peroxisome-targeted mCherry, while co-localization with mitochondrial-targeted mCherry was not apparent (Fig 7A). Immunofluorescence was used as a complementary approach. Fluorescence from the anti-TMEM135 antibody was low in HepG2 cells. To improve the signal to noise ratio, cells were transfected with a plasmid encoding endogenous TMEM135. Peroxisomes and mitochondria were visualized with mouse monoclonal antibodies against ABCD3 and COX4I1, respectively. Using this method the TMEM135 antibody revealed punctate cytoplasmic structures that co-localized with the ABCD3 antibody, but did not appear to co-localize with the COX4I1 antibody (Fig 7B).

**Fig 7.**
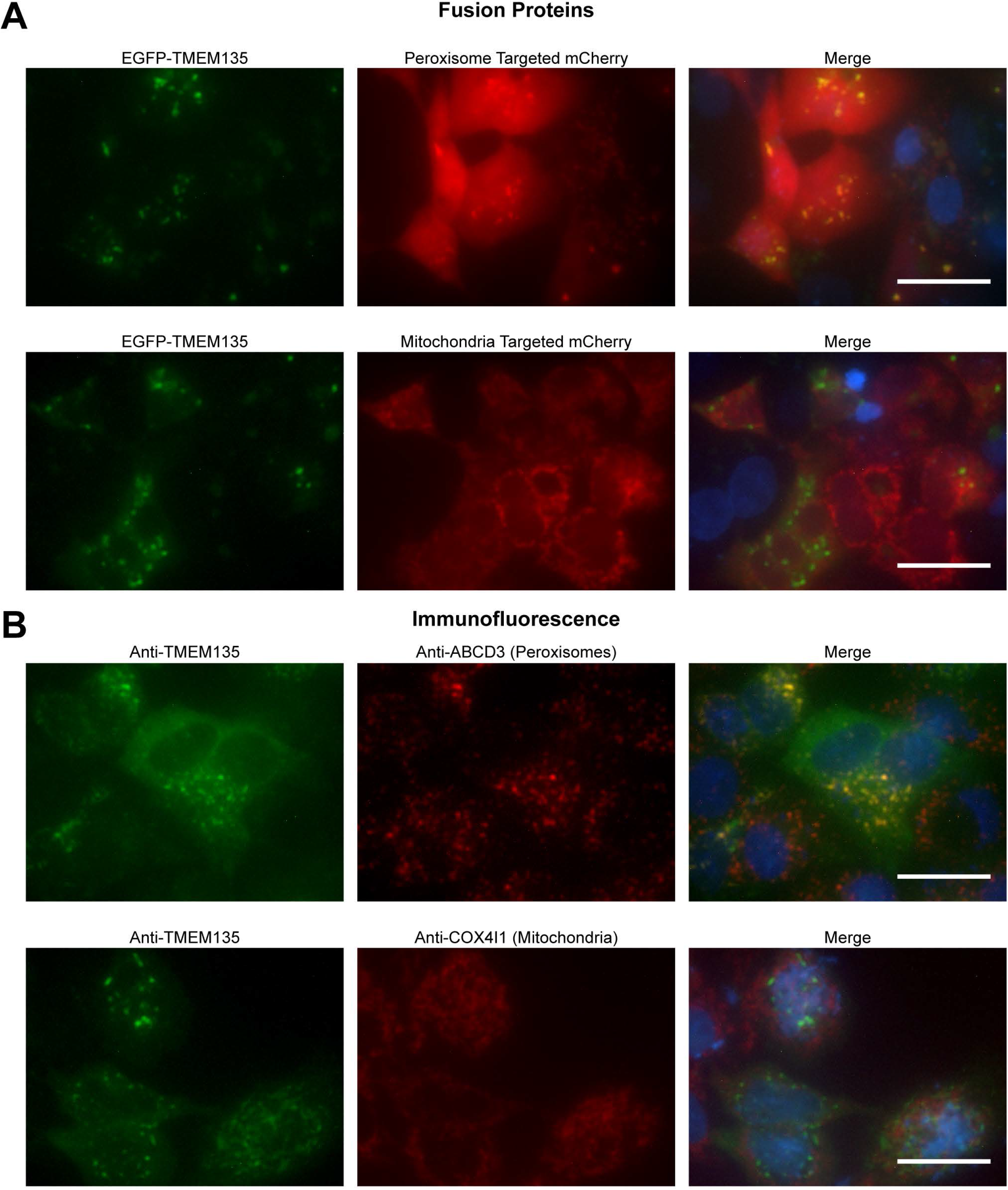
TMEM135 is localized to peroxisomes while mitochondrial localization is not apparent. Panel A is images of HepG2 cells transiently expressing an EGFP-TMEM135 fusion protein and peroxisomal or mitochondrial targeted mCherry. Panel B is immunofluorescence localization of TMEM135 and peroxisomes or mitochondria. HepG2 cells were transiently transfected with the pTarget-*TMEM135* plasmid to increase detection sensitivity in immunofluorescence. For both panels, nuclei are shown in blue in the merged images, and scale bars represent 25 μm.

### TMEM135 knockdown inhibits fatty acid metabolism during fasting

We next determined the effect of TMEM135 on markers of peroxisomal metabolism. Because fasting causes lipolysis in adipose and increased fatty acid uptake and β-oxidation in the liver, we determined hepatic fatty acid concentrations. As expected, fasting induced a significant increase in hepatic concentrations of several fatty acids in mice receiving the control siRNA (Table 2). In fasted mice, TMEM135 knockdown resulted in a further significant increase in total fatty acids and linoleic acid (Fig 8), with trends (0.05 < p < 0.1) for increases in several other fatty acids (Table 2). In general, mitochondria preferentially oxidize short and medium chain fatty acids (<C12), both mitochondria and peroxisomes oxidize long chain fatty acids (LCFA, C14-C18), while peroxisomes exclusively oxidize very-long chain fatty acids (VLCFA, >C20) [38]. While mitochondria are believed to be the principal site of LCFA β-oxidation, peroxisomes also directly oxidize LCFA in a cooperative manner with mitochondria [39]. The peroxisomal contribution to LCFA β-oxidation becomes quantitatively greater during physiologic states of increased fatty acid load such as fasting [34]. The significant increase in linoleic and total fatty acids is consistent with an impairment in β-oxidation in fasted TMEM135 knockdown mice, and when considering hepatic NADH and ketone concentrations (Fig 5E-F) and TMEM135 localization (Fig 7), indicates that the impairment occurred in peroxisomes. Also, linoleic acid is an essential fatty acid obtained via dietary sources [40], so alterations in fatty acid synthesis or desaturation cannot explain its increased concentrations. Interestingly, in fed mice TMEM135 knockdown significantly decreased 22:0 and tended to decrease 24:0, indicating an increase in β-oxidation of VLCFA in the fed state. Peroxisomal substrate preference can be modified as bezafibrate treatment increases LCFA but not VLCFA β-oxidation in rat liver peroxisomes [41]. Fasting results in PPARA activation similar to bezafibrate treatment, and our data are consistent with TMEM135 contributing to a switch in peroxisomal preference for LCFA compared to VLCFA. Regarding bile acids, hepatic concentrations of cholic acid were not significantly altered by TMEM135 knockdown (Fig S6). Collectively, these data indicate that TMEM135 may not be obligatory to basal peroxisome function, but under physiologic conditions of elevated hepatic fatty acid flux such as fasting, TMEM135 may play a key role in enhancing peroxisomal metabolism.

**Fig 8.**
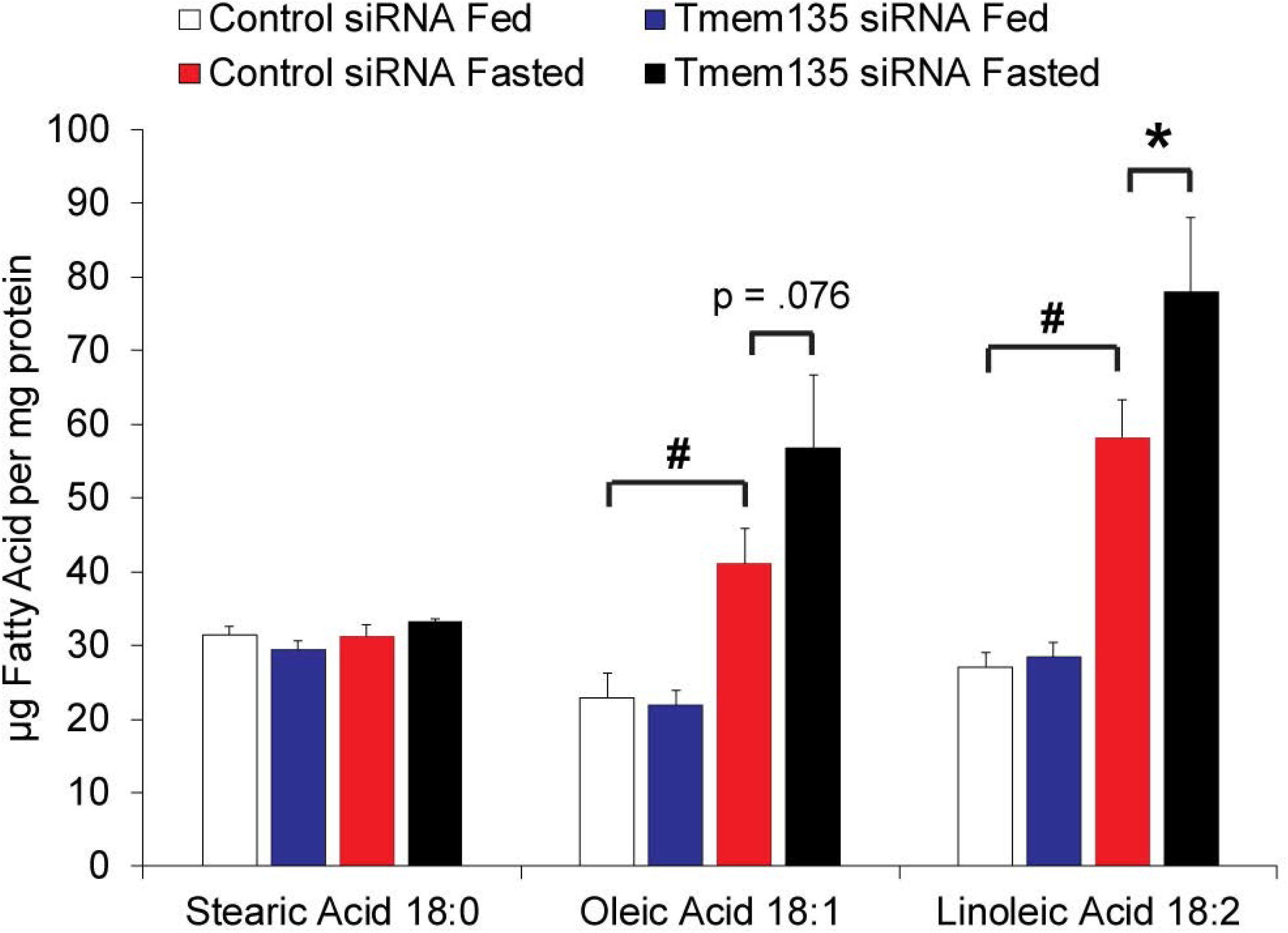
TMEM135 mediates fasting-induced fatty acid metabolism. The concentrations of C18 fatty acids quantified by GC are plotted. Asterisk denotes significant difference (p< 0.05) within feeding status due to *Tmem135* siRNA, and # indicates significant difference due to fasting in mice receiving the control siRNA (n = 5 per siRNA and nutritional status).

**Table 2.**
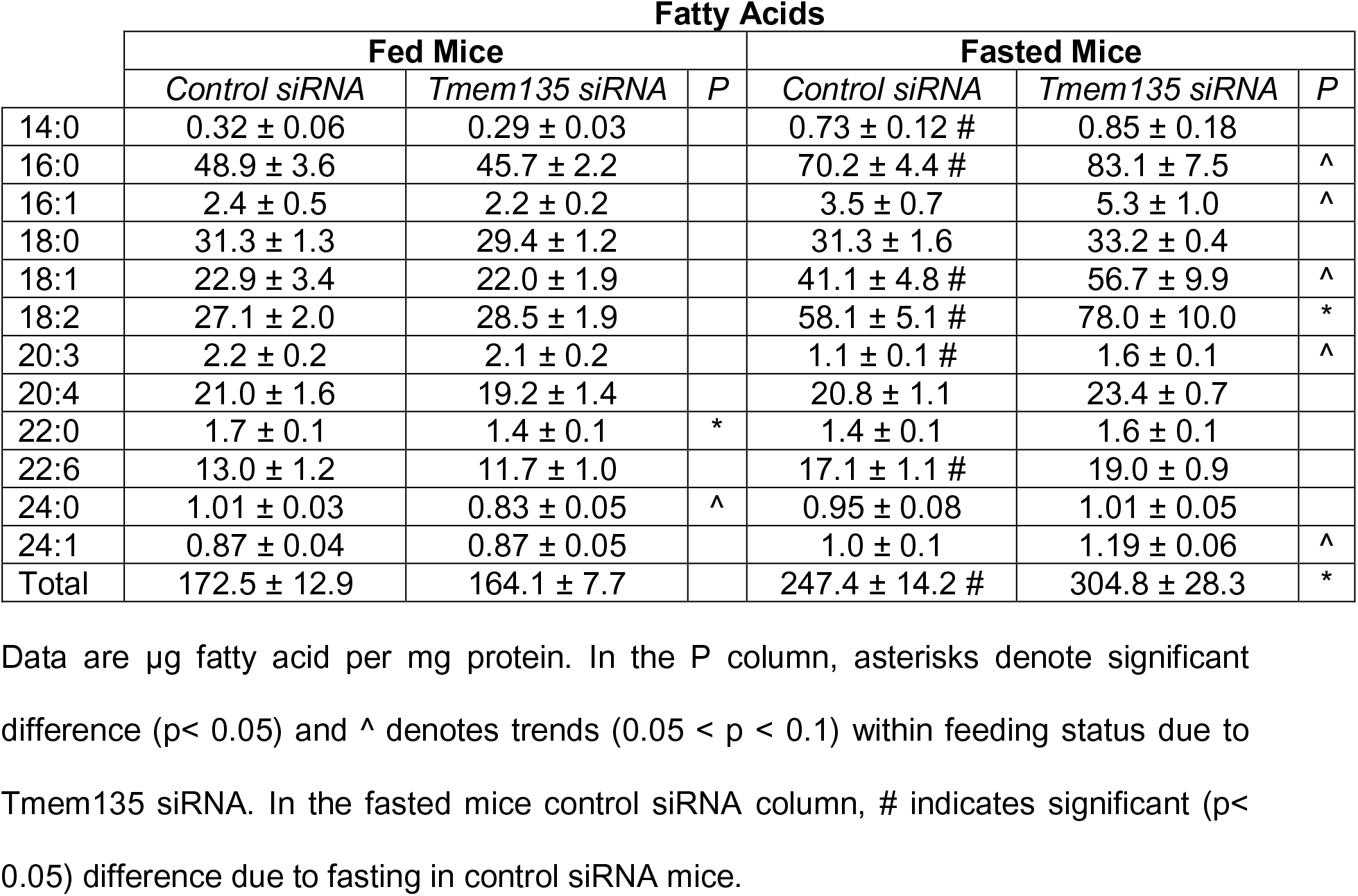
Fatty acids quantified by GC

## Discussion

In the current study we determined that TMEM135 is an LXR-inducible protein in humans that controls peroxisomal metabolism by regulating matrix enzyme concentrations. The regulation of TMEM135 by the LXRs in humans (and nonhuman primates) is supported by several observations including: 1) the LXR agonist T09 stimulates TMEM135 mRNA and protein in multiple immortalized human cell lines, as well as in primary rhesus macaque luteal cells, 2) LXR knockdown prevents T09-induced *TMEM135* mRNA expression, 3) EMSA and ChIP analysis demonstrated LXR binding to an LXRE in the *TMEM135* promoter, and 4) mutation of the LXRE within the *TMEM135* promoter prevents LXR agonist-induced expression. A mouse model was used to determine the *in vivo* importance of TMEM135 because it is the most conducive to genetic manipulation. Because mouse *Tmem135* is not an LXR target gene, it is not possible to test the effect of the LXRs on TMEM135-mediated peroxisomal metabolism in this model. However, given that we have identified the functional LXRE in the human *TMEM135* promoter, a humanized mouse line could be created for future studies to test LXR regulation of TMEM135 *in vivo*. Physiologic data was generated using a liver-specific, partial knockdown of TMEM135. Obviously, a genetic knockout would be preferable. Given the limited information on TMEM135 function and the cost and time required to create knockout mice, siRNA was used as a first-line approach to determine its physiologic importance. It is reasonable to predict that the *in vivo* effects of partial TMEM135 knockdown observed in the current study will be amplified in knockout mice. Based on the findings from the current study, we have begun developing TMEM135 knockout mice for future studies.

Multiple observations in mice collectively support TMEM135 as being a regulator of peroxisomal metabolism. When hepatic β-oxidation (both peroxisomal and mitochondrial) was stimulated by fasting, TMEM135 knockdown mice had significantly lower NADH (Fig 5E) and significantly higher total fatty acid and linoleic acid concentrations (Fig 8 and Table 2), consistent with an impairment in β-oxidation. Meanwhile, hepatic ketones tended to increase in knockdown mice when fasted (Fig 5F), and ketogenesis occurs exclusively in mitochondria and is intricately linked to β-oxidation [31]. Taken together, these findings indicate that β-oxidation was impaired in peroxisomes, but not in mitochondria. This is further supported by our finding that TMEM135 localizes to peroxisomes (Fig 7). The increase in hepatic fatty acid concentrations in fasted mice (Fig 8 and Table 2) parallels the increase in triglycerides that occurred in HepG2 cells (Fig 4) following TMEM135 knockdown, indicating that TMEM135 likely plays a similar role in mouse and human cells. Mechanistically, TMEM135 may modulate peroxisomal metabolism by regulating matrix enzyme concentrations within peroxisomes consistent with its homology to the Tim17 family of protein translocases [32], although additional effects on peroxisome number or stability are possible.

Direct assays of β-oxidation were not performed in the current study because they are unlikely to further delineate peroxisomal from mitochondrial β-oxidation. Fatty acid quantification shown in Fig 8 and Table 2 indicates that TMEM135 may help stimulate LCFA metabolism, but not VLCFA metabolism. As both mitochondria and peroxisomes can utilize LCFA as substrates [39], β-oxidation assays using LCFA would not further distinguish between mitochondrial and peroxisomal contributions unless highly purified peroxisome fractions could be attained. However, these were not obtained in the current study because when tissue collection occurred we were working under the assumption that TMEM135 was a mitochondrial protein based on previous reports of its association with mitochondria [26,33]. Livers were frozen, which precludes the isolation of highly purified peroxisome fractions that require fresh tissue. Furthermore, a CPT1A inhibitor such as etomoxir cannot be used to differentiate mitochondrial from peroxisomal β-oxidation because peroxisomal β-oxidation produces chain-shortened fatty acids that can enter the mitochondria in a CPT1A-independent manner [39]. Future studies in genetic TMEM135 knockout mice should be used to directly assess the role of TMEM135 in peroxisomal β-oxidation.

The LXRs are master regulators of reverse cholesterol transport (RCT), a process whereby excess cholesterol is removed from peripheral tissues and transported to the liver for elimination in bile acids [1]. There has been much interest in the development of LXR agonists to treat diseases including atherosclerosis, Alzheimer’s disease, and other metabolic disorders [1]. However, the therapeutic potential of LXR agonists has been limited because they also increase hepatic lipogenesis via induction of SREBP1c [8,9]. Our data indicate that the LXRs can stimulate peroxisomal β-oxidation by increasing transcription of *TMEM135*, which consequently may limit steatosis during LXR-induced lipogenesis. It should be noted that because *TMEM135* is an LXR target gene in humans and not mice, LXR agonist-induced steatosis may be less pronounced in human compared to mice hepatocytes due to species differences in LXR regulation of *TMEM135* transcription. A mouse line with the human LXRE inserted into the *Tmem135* promoter could address this. Previous evidence of a link between the LXRs and peroxisomal β-oxidation is limited and conflicting. One group reported increased hepatic peroxisomal β-oxidation in mice treated with an LXR agonist through an undefined mechanism involving increased oxidative gene expression [42,43]. However, another report indicates that the LXRs suppress peroxisomal β-oxidation by inhibiting expression of the peroxisomal membrane transporter ABCD2 [44]. It would be interesting to determine how LXR-mediated suppression of ABCD2 and induction of TMEM135 influences net peroxisome function.

Previous studies to determine the biological function of TMEM135 are limited. The first indication of its physiologic role came from a mouse model of mitochondrial acyl-CoA dehydrogenase very long chain (ACADVL) deficiency [26]. Under conditions of increased demand for β-oxidation such as fasting and cold stress, ACADVL-deficient mice have reduced survival due to cardiac dysfunction [26]. It was discovered that TMEM135 was elevated more than four-fold in the hearts of ACADVL-deficient mice that survived birth [26]. The ACADVL enzyme catalyzes the first step in mitochondrial β-oxidation using fatty acids larger than 14 carbons as substrates [45]. Because ACADVL deficiency restricts the mitochondria from directly oxidizing LCFA, data from the current study indicate that the increase in TMEM135 may have promoted survival by increasing peroxisomal metabolism of LCFA to medium chain fatty acids that were subsequently oxidized in the mitochondria in an ACADVL-independent manner. Another study reported that a mutation in *Tmem135* was responsible for accelerated aging of the retina in a mouse model of age-related macular degeneration [33], and neurodegenerative disorders are commonly associated with peroxisome dysfunction [38]. Both of these studies on TMEM135 reported that it is localized to mitochondria [26,33], which conflicts with findings in the current study that it is a peroxisomal protein. The current findings are consistent with two earlier studies that originally detected TMEM135 (referred to as peroxisome membrane protein 52 or PMP52) via proteomics in highly purified peroxisome fractions from rodents, and further confirmed its localization to peroxisomes but not mitochondria [41,46]. Neither of the studies that reported TMEM135 localization to mitochondria tested peroxisomal localization and they both described TMEM135 in punctate structures that were near, but not necessarily within, mitochondria [26,33]. Because peroxisomes and mitochondria can physically associate via the formation of tethering complexes [47], it seems likely that reports of TMEM135 association with mitochondria [26,33] were due to peroxisomes that were tethered to mitochondria, and TMEM135 effects on mitochondria are indirectly mediated via altered peroxisomal metabolism. Collectively, previously published and current data indicate that TMEM135 is a peroxisomal and not a mitochondrial protein. Additional studies are needed to more precisely determine the indirect effects of TMEM135 on mitochondrial function.

The critical role of peroxisomes in human health is illustrated by the spectrum of peroxisome disorders; but in addition to these inborn errors of metabolism it is becoming increasingly apparent that peroxisomes play key roles in other metabolic and age-related diseases including cancer, neurodegenerative disorders, and diabetes [38]. This may be due to direct catabolic and anabolic actions of peroxisomes, as well as interactions between peroxisomes and other organelles including mitochondria [38,47]. Our data indicate that TMEM135 regulates peroxisomal metabolism by increasing matrix enzyme concentrations. Furthermore, the effects of TMEM135 are inducible as it is an LXR target gene in humans, and it may be regulated by other metabolic signaling pathways. In support of this, PPARA is the master regulator of β-oxidation in the liver and in primary human hepatocytes *TMEM135* mRNA expression is consistently induced by PPARA agonists [27] and its promoter is bound by PPARA as determined using ChIP-seq [48], indicating it is also a PPARA target gene. These findings implicate TMEM135 as a potential therapeutic target in the treatment of metabolic and age-related diseases that are associated with peroxisome dysfunction.

## Acknowledgements

Mass spectrometry and proteomics data were acquired by the Arizona Proteomics Consortium at the BIO5 Institute of the University of Arizona. Microarray data were acquired by the Oregon Health & Science University Affymetrix Microarray Core. The authors would like to thank Dr. André-Denis Wright for arranging Departmental support to ensure completion of this project.

