## Supplementary material for "TMEM135 is an LXR-inducible regulator of peroxisomal metabolism": Fig S

### Supporting Information

**S1 File. Excel file of filtered microarray data.**

**S1 Table. Primer sequences and detection method used in QPCR.**

**S1 Figure. Relative locations of potential LXREs in the human *TMEM135* promoter.** The relative location of each potential LXRE is indicated with a number in the box to identify it (1 = LXRE1, 2 = LXRE2, 3 = LXRE3). Transcription and translation start sites of *TMEM135* based on NCBI accession numbers NM_022918.3 (TSS1) and NM_022918.4 (TSS2), with the latter replacing the former on November 23, 2018. In our data, transcripts corresponding to both accession numbers were expressed (see Fig. 3). Arrows indicate the putative transcriptional start sites (TSS) for each respective transcript, with ATG identifying the common translational start site. Numbers above each LXRE indicate the number of base pairs that the first nucleotide at the 5' end of the LXRE is from the common translational start site (+1). Panel B is sequences of each LXRE and mutated LXRE generated for EMSA and reporter assays. The hexanucleotide half-sites are indicated by capitalized bold letters, mutated nucleotides are underlined.

**S2 Figure. *Tmem135* is not an LXR target gene in mice.** Panel A compares the LXRE3 sequence between humans and mice. The hexanucleotide half-sites of the DR-4 sequence are shown in capitalized bold letters. A one nucleotide deletion in the mouse sequence relative to humans is indicated by a double dash. Panels B and C display the effect of a 24-hour T09 treatment on *Tmem135* and *Abca1* mRNA expression normalized to *Mrps10*, respectively, in immortalized mouse hepatocyte (BNL 1NG A.2) and macrophage (RAW 264.7) cell lines (n = 4). Asterisks denote significant increase ($p < 0.05$) in mRNA expression due to T09.

**S3 Figure. TMEM135 mediates proliferation of HepG2 cells.** Panel A is the effect of TMEM135 knockdown on proliferation of HepG2 cells. Asterisks denote significant difference at the corresponding timepoint (n = 4). Panel B is the effect of TMEM135 knockdown on cell cycle progression in HepG2 cells, and panel C is the effect on mRNA expression of cell cycleassociated genes normalized to *MRPS10* and expressed as fold-change relative to the control siRNA. For panels B-C, asterisks denote significant differences due to TMEM135 knockdown (n = 4). Panel D is the effect of TMEM135 knockdown on ATP content (normalized to total protein content) in HepG2 cells after a 4-hour incubation in glucose-free, sodium pyruvate-free media. The chart is plotted with the control siRNA set to 1, and the asterisk denotes a significant difference (n = 4).

**S4 Figure. Effect of *in vivo* TMEM135 knockdown on hepatic metabolic markers.** Panel A is liver triglyceride concentrations, panel B is liver non-esterified fatty acid (NEFA) concentrations, panel C is liver ATP concentrations, and panel D is liver glycogen concentrations. For all panels, # indicates significant difference ($p < 0.05$) due to fasting in animals receiving the control siRNA (n = 5).

**S5 Figure. Effect of *in vivo* TMEM135 knockdown on serum metabolic markers.** Panel A displays lipids, panel B is NEFAs, panel C is glucose, panel D is insulin, and panel E is β-hydroxybutyrate in serum. For all panels, # indicates significant difference ($p < 0.05$) due to fasting in animals receiving the control siRNA (n = 5).

**S6 Figure. Effect of *in vivo* TMEM135 knockdown on hepatic cholic acid concentrations.** Concentrations of the primary bile acid, cholic acid, are plotted. # indicates significant difference ($p < 0.05$) due to fasting in animals receiving the control siRNA (n = 5).

**Table S1.** Primer sequences and detection method used in QPCR.

| | | | |
|---|---|---|---|
| | ***Human mRNA Targets*** | | |
| **Gene** | **Forward primer (5'-3')** | **Reverse primer (5'-3')** | **MGB Probe (5'-3') or SYBR** |
| *ABCA1* | TCCAGGCCAGTACGGAATTC | TCCTCGCCAAACCAGTAGGA | CTGGTATTTTCCTTGCACCAA |
| *MRPS10* | TTCCAAAGGATTTGACCAAACC | TCGTGACCTTTCACCAAAACC | ATCTCTGATGAACCAGACAT |
| *NR1H2* | ATCGTGGACTTCGCTAAGCAA | GATCTCGATAGTGGATGCCTTCA | TGCCTGGTTTCCTGC |
| *NR1H3* | TCCCCATGACCGACTGATG | CAGACGCAGTGCAAACACTTG | TCCCACGGATGCTAAT |
| *TMEM135* | CATGAGGAAAAACCCGGAAGA | ACATCTATGCCTTGGTCCATGTT | TTGAATCCACAAATTTCCT |
| *TMEM135 LXRE3* | CGCTCAACATCCGAGGACTT | ACTCGGAGGTGCGGAATG | CCCCGCTCTCGTGAC |
| *ChIP Non-Specific Locus* | GGCTCTTGTAGGAGTGATGTCACA | CAACTGCCAGAGGGACACTTG | CATGTCCTTGGTATGCCT |
| *PPARA* | GAAGGAACTTCGGTTGTGTAAAGG | CCAAGACGTGCCCAATGTC | SYBR Green |
| *CPT1A* | CGGGAGGAAATCAAACCAATT | GGGATCCGGGAAGTATTAAACAT | SYBR Green |
| *SREBF1* | GGAGCCATGGATTGCACTTT | AGCATAGGGTGGGTCAAATAGG | SYBR Green |
| *FASN* | GCAAATTCGACCTTTCTCAGAAC | CTCGTTGAAGAACGCATCCA | SYBR Green |
| *CDKN2A* | CAGTAACCATGCCCGCATAGA | AAGTTTCCCGAGGTTTCTCAGA | SYBR Green |
| *TP53* | AGTGTGGTGGTGCCCTATGAG | GCCCATGCAGGAACTGTTACA | SYBR Green |
| *CDK2* | CTCTGCTCTCACTGGCATTCC | ACCCGATGAGAATGGCAGAA | SYBR Green |
| *CCNE1* | AGCCAGCCTTGGGACAATAA | TGGGTAAACCCGGTCATCAT | SYBR Green |
| *CCNA2* | CCTGCGTTCACCATTCATGT | CAGGGCATCTTCACGCTCTAT | SYBR Green |
| | ***Mouse mRNA Targets*** | | |
| *Tmem135* | CATGGACCAAGGCACAGATG | CAAGTAGCCCACGCTGAACA | SYBR Green |
| *Mrps10* | CCAGCGAAACTTGCCTGAAG | TCCCACATCGGTTCCTTGAT | SYBR Green |
| *Ppara* | AGAGCCCCATCTGTCCTCTC | ACTGGTAGTCTTGCAAAACCAAA | SYBR Green |
| *Cpt1a* | CCTGGGCATGATTGCAAAG | AAGAGGACGCCACTCACGAT | SYBR Green |
| *Acadm* | AACACAACACTCGAAAGCGG | TTCTGCTGTTCCGTCAACTCA | SYBR Green |
| *Ucp2* | ATGGTTGGTTTCAAGGCCACA | CGGTATCCAGAGGGAAAGTGAT | SYBR Green |
| *Sirt3* | ACAGGCCCAATGTCACTCACT | CAAGCCCGTCGATGTTCTG | SYBR Green |
| *Abca1* | GGACATGCACAAGGTCCTGA | CAGAAAATCCTGGAGCTTCAAA | SYBR Green |

**A**

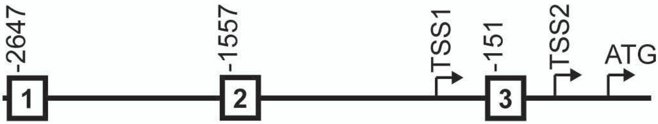


**B**

**WT LXRE1**
5-tggt**GGATCA**cctg**AGGTCA**taag-3
3-acca**CCTAGT**ggac**TCCAGT**attc-5

**MUT LXRE1**
5-tggt**TAATCA**cctg**ACATCA**taag-3
3-acca**ATTAGT**ggac**TGTAGT**attc-5

**WT LXRE2**
5-gggt**GGATCA**cttg**AGGTCA**ggag-3
3-ccca**CCTAGT**gaac**TCCAGT**cctc-5

**MUT LXRE2**
5-gggt**TAATCA**cttg**ACATCA**ggag-3
3-ccca**ATTAGT**gaac**TGTAGT**cctc-5

**WT LXRE3**
5-agcg**GGGTTA**ctct**GGGCCA**aagt-3
3-tcgc**CCCAAT**gaga**CCCGGT**ttca-5

**MUT LXRE3**
5-agcg**TAGTTA**ctct**ACACCA**aagt-3
3-tcgc**ATCAAT**gaga**TGTGGT**ttca-5

Figure S1

**A**

**Human** 5-agcg**GGGTTA**ctct**GGGCCA**aagt-3

**Murine** 5-agca**GG--TTA**ctca**AGGCCG**gagt-3

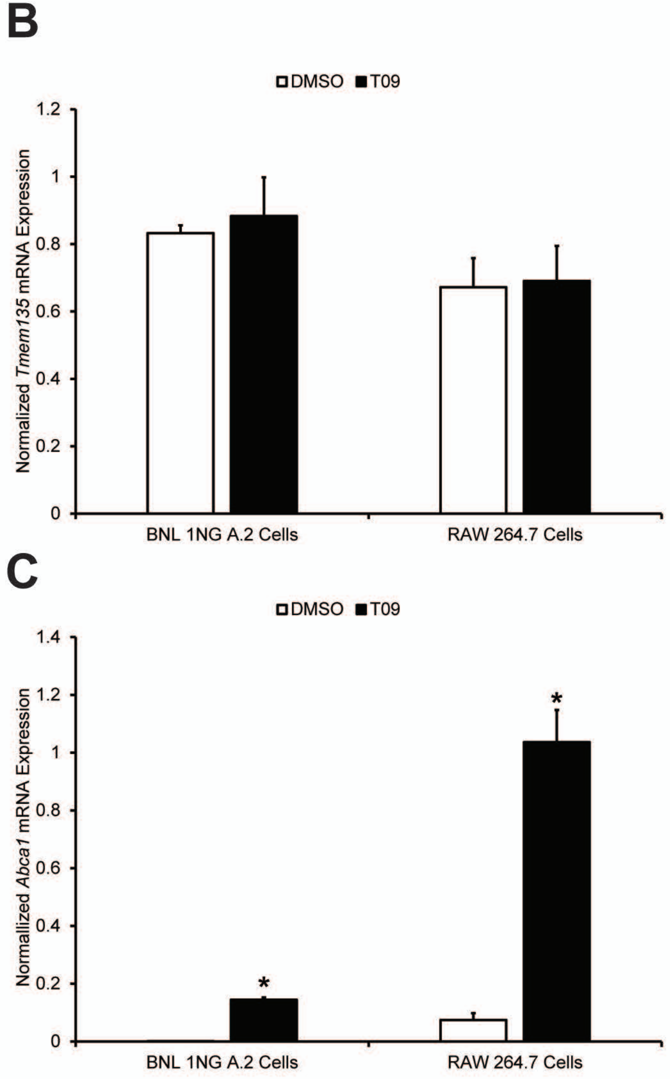


Figure S2

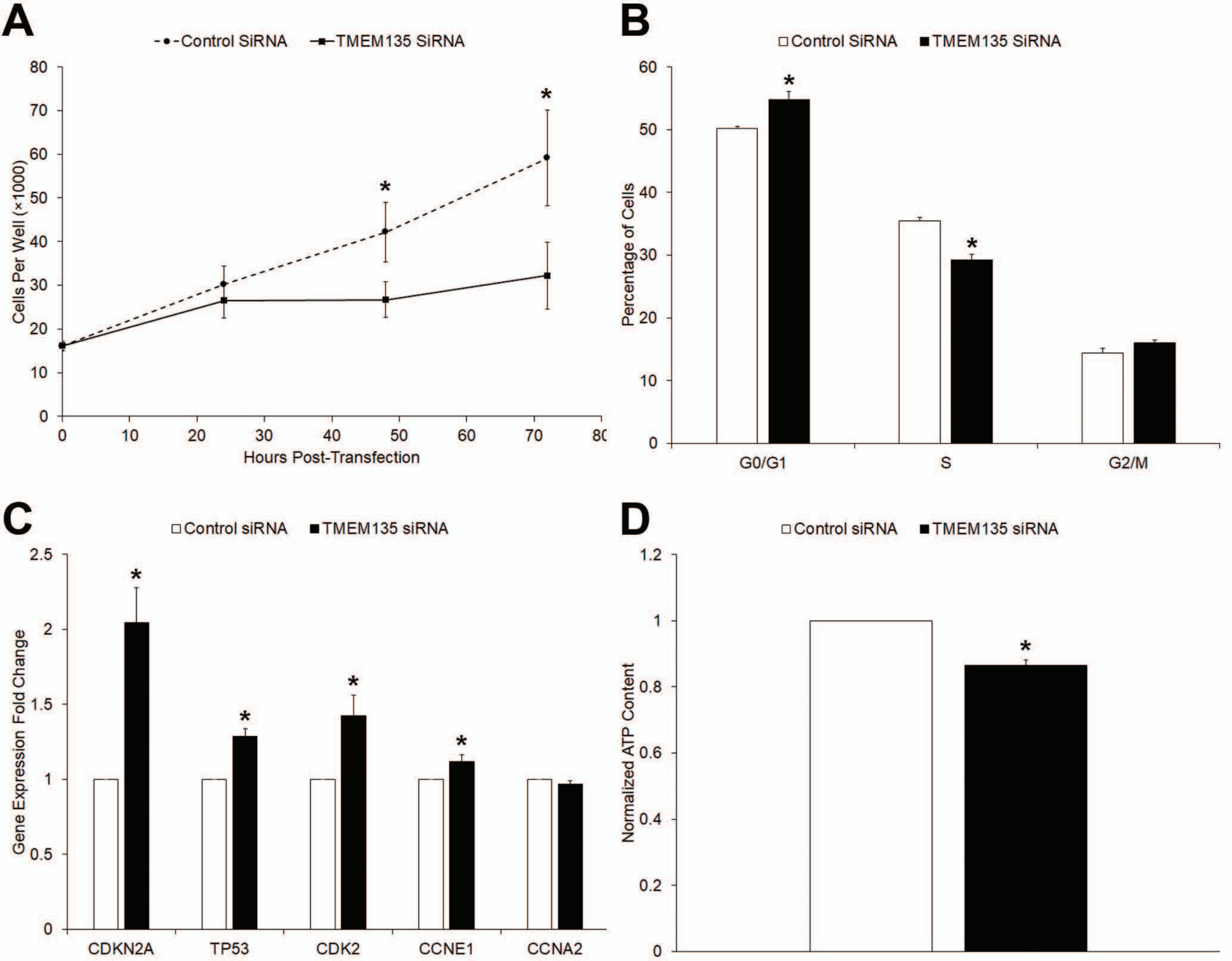
A
Control SiRNA
TMEM135 SiRNA
Cells Per Well (×1000)
Hours Post-Transfection
B
Control SiRNA
TMEM135 SiRNA
Percentage of Cells
G0/G1
S
G2/M
C
Control siRNA
TMEM135 siRNA
Gene Expression Fold Change
CDKN2A
TP53
CDK2
CCNE1
CCNA2
D
Control siRNA
TMEM135 siRNA
Normalized ATP Content

Figure S3

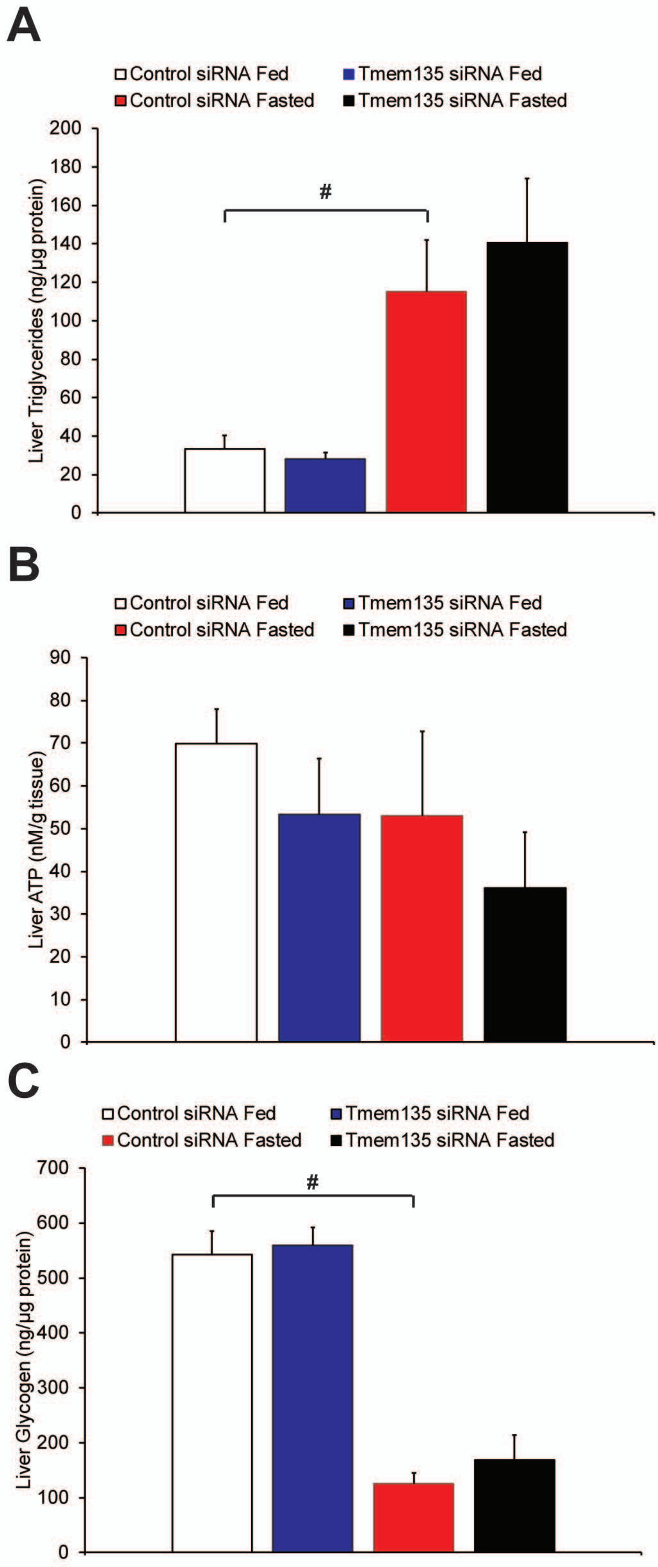
A
Control siRNA Fed
Tmem135 siRNA Fed
Control siRNA Fasted
Tmem135 siRNA Fasted
Liver Triglycerides (ng/µg protein)
0
20
40
60
80
100
120
140
160
180
200
#
B
Control siRNA Fed
Tmem135 siRNA Fed
Control siRNA Fasted
Tmem135 siRNA Fasted
Liver ATP (nM/g tissue)
0
10
20
30
40
50
60
70
80
90
C
Control siRNA Fed
Tmem135 siRNA Fed
Control siRNA Fasted
Tmem135 siRNA Fasted
Liver Glycogen (ng/µg protein)
0
100
200
300
400
500
600
700
#

Figure S4

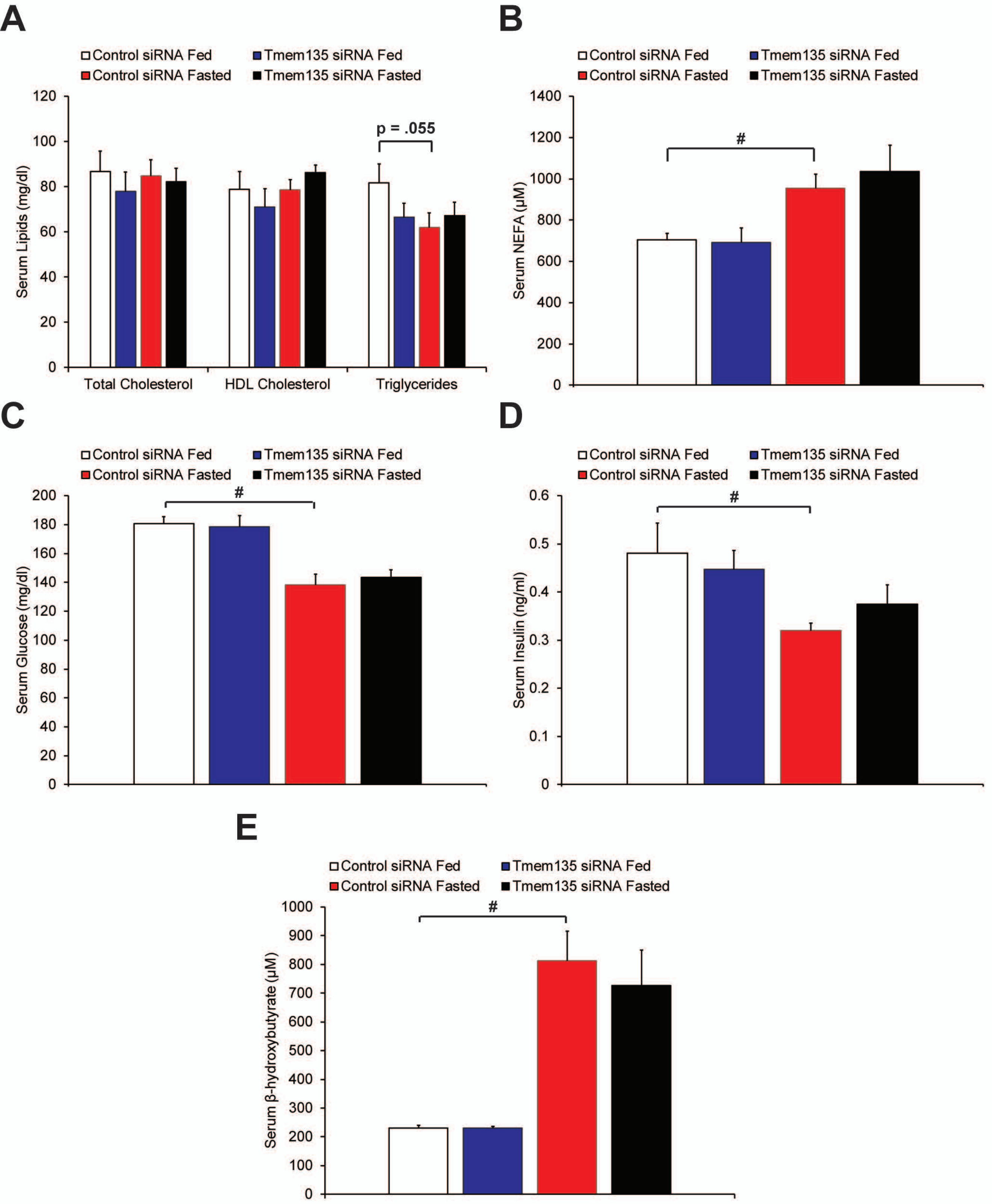

A
Control siRNA Fed
Tmem135 siRNA Fed
Control siRNA Fasted
Tmem135 siRNA Fasted
Serum Lipids (mg/dl)
p = .055
Total Cholesterol
HDL Cholesterol
Triglycerides
B
Serum NEFA (μM)
#
C
Serum Glucose (mg/dl)
#
D
Serum Insulin (ng/ml)
#
E
Serum β-hydroxybutyrate (μM)
#


Figure S5

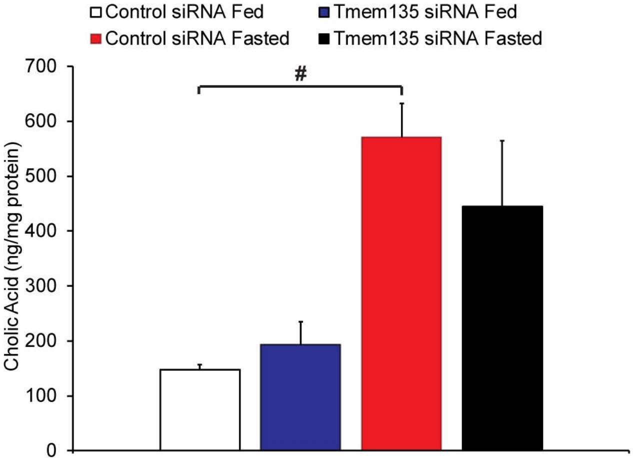

Control siRNA Fed
Tmem135 siRNA Fed
Control siRNA Fasted
Tmem135 siRNA Fasted
700
600
500
400
300
200
100
0
Cholic Acid (ng/mg protein)
#


Figure S6
